# Parallel nonfunctionalization of CK1δ/ε kinase ohnologs following a whole-genome duplication event

**DOI:** 10.1101/2023.10.02.560513

**Authors:** Daniel Evans-Yamamoto, Alexandre K Dubé, Gourav Saha, Samuel Plante, David Bradley, Isabelle Gagnon-Arsenault, Christian R Landry

## Abstract

Whole genome duplication (WGD) followed by speciation allows us to examine the parallel evolution of ohnolog pairs. In the yeast family *Saccharomycetaceae*, *HRR25* is a rare case of repeated ohnolog maintenance. This gene has reverted to a single copy in *S. cerevisiae* where it is now essential, but has been maintained as pairs in at least 7 species post WGD. In *S. cerevisiae*, *HRR25* encodes the casein kinase (CK) 1δ/ε and plays a role in a variety of functions through its kinase activity and protein-protein interactions (PPIs). We hypothesized that the maintenance of duplicated *HRR25* ohnologs could be a result of repeated subfunctionalization. We tested this hypothesis through a functional complementation assay in *S. cerevisiae*, testing all pairwise combinations of 25 orthologs (including 7 ohnolog pairs). Contrary to our expectations, we observed no cases of pair-dependent complementation, which would have supported the subfunctionalization hypothesis. Instead, most post-WGD species have one ohnolog that failed to complement, suggesting their nonfunctionalization or neofunctionalization. The ohnologs incapable of complementation have undergone more rapid protein evolution, lost most PPIs that were observed for their functional counterparts and singletons from post and non-WGD species, and have non-conserved cellular localization, consistent with their ongoing loss of function. The analysis in *N. castelli* shows that the non-complementing ohnolog is expressed at a lower level and has become non-essential. Taken together, our results indicate that *HRR25* orthologs are undergoing gradual nonfunctionalization.

## Introduction

One of the driving forces behind the evolution of complex traits and the diversification of species is gene duplication (Otto and Whitton 2000; Donoghue and Purnell 2005). Although most genes are duplicated at one point or another during their evolutionary history, only a fraction will be retained in some lineages as duplicated pairs or as part of larger families. Identifying the molecular mechanisms and population dynamics parameters that dictate the retention of some gene duplicates, but not others, is a major goal in molecular evolution.

The classical model for the maintenance of gene duplication posits that due to its redundancy, a duplicated gene is now free to explore a new sequence space and therefore can gain new, adaptive functions that favor its maintenance (Ohno 1970) (**Figure 1 A**). Another, alternative model is that since genes have multiple functions and loss of function through mutations is generally more likely than gain of function, duplicated genes could simply partition the ancestral function and functionally complement each other, resulting in two genes that together only can perform the ancestral function (subfunctionalization) (Hughes 1994; Force et al. 1999). In this model, the paralogs degenerate and eventually complement each other (DDC; Duplication, Degeneration and Complementation). These models apply not only to protein functions but also to their expression. For instance, gene duplication could simply increase protein dosage and that in itself could be adaptive, leading to the maintenance of the two genes with no necessary changes in the intrinsic molecular functions of the proteins (Innan and Kondrashov 2010; Kondrashov 2010). Indeed, increases in gene copy number have been shown to be adaptive in many cases (Perry et al. 2007; Pajic et al. 2019; Ascencio et al. 2021). When increased expression is not adaptive, subfunctionalization could occur at the level of expression alone. For instance, if duplicates evolve to have lower expression levels after gene duplication, the paralogs as a pair are required to achieve the dosage that is needed to perform the ancestral function (Qian et al. 2010; Gout and Lynch 2015). In this case again, no divergence of function is required. However, once divergence in expression becomes asymmetrical, the asymmetry could enhance with time and the least expressed paralog could get to a level of expression low enough that further degenerative mutations affecting expression, function, or stability would become effectively neutral. Therefore, although initially the genes are maintainted by dosage subfunctionalization, this mechanism could eventually promote slow gene loss in the long term (Gout and Lynch 2015). Gene loss through nonfunctionalization eventually occurs via this model but not as rapidly as it would have if loss-of-function mutations occurred soon after gene duplication.

**Figure 1.**
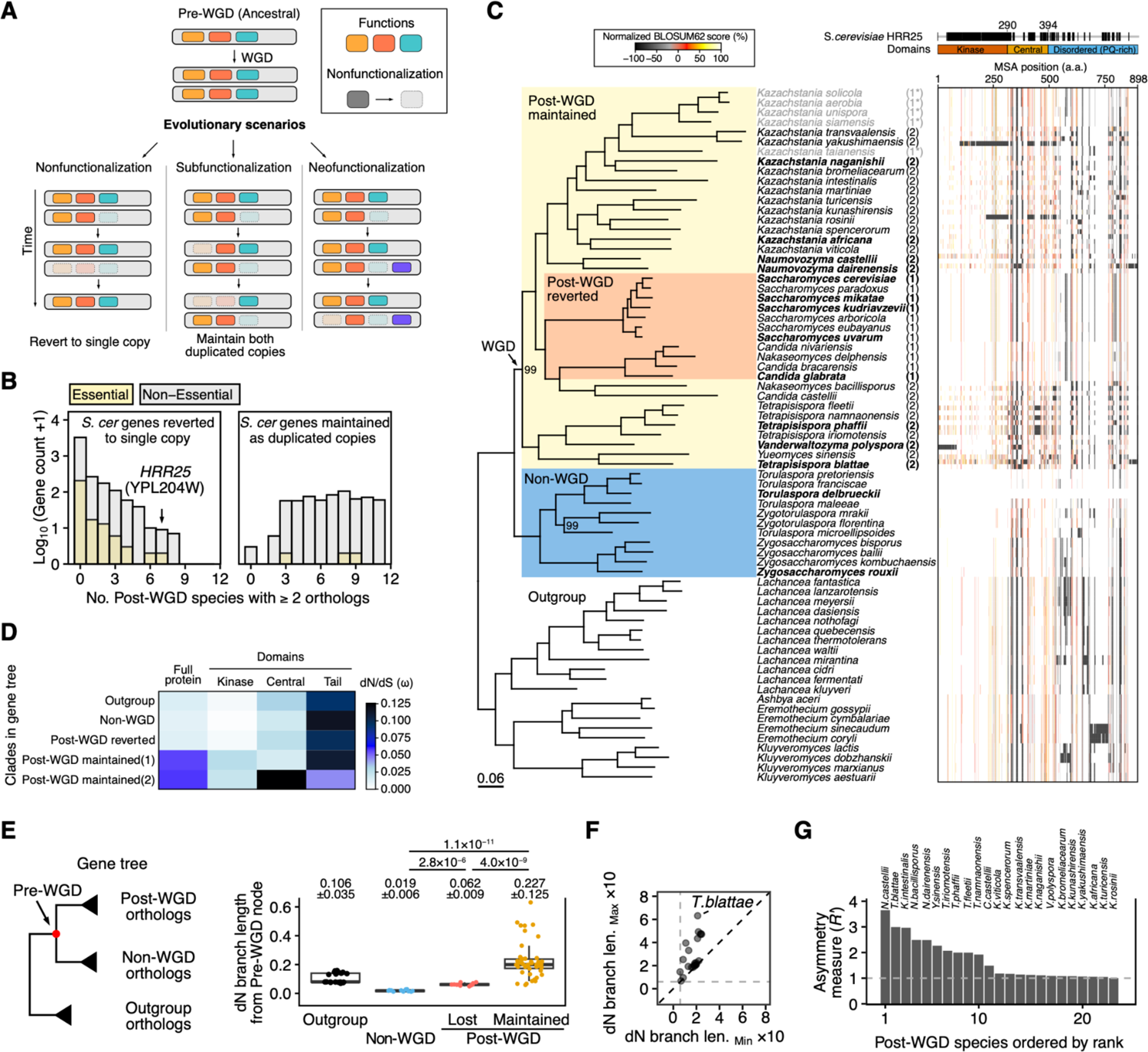
Conservation and divergence of *HRR25* ohnologs in the *Saccharomycetaceae* family after the whole-genome duplication. **(A)** Diagram of evolutionary fates of ohnologs after WGD. (Left) One of the ohnologs accumulates mutations and nonfunctionalizes. (Middle) Subfunctionalization: two copies are partially degenerating, each preserving parts of the ancestral functions of the protein. (Right) Neofunctionalization: one copy maintains the ancestral function and another gains novel functions. (**B**) Number of post-WGD species that maintained both ohnologs for essential and non essential *S. cerevisiae* genes and separated by status in *S.cerevisiae*. *HRR25* is an exceptional case of ohnologs that were maintained in several species while returning to a single copy in *S. cerevisiae*. **(B)** (Left) Phylogenetic tree of *Saccharomycetaceae*, adapted from (Li et al., 2021). Confidence annotations for branches with bootstrap confidence of 100 are abbreviated. Post-WGD species are annotated with respect to the numbers of *HRR25* copies. Species used for homology search are shown in bold. Species with low confidence for copy number annotation are in light gray and number is marked with an asterisk. (Right) Multiple sequence alignment of Hrr25p. Normalized BLOSUM62 score for divergence with the non-WGD *T.delbrueckii* ortholog (TDEL0E05230). The aligned region of *S.cerevisiae* Hrr25p is indicated on top as black boxes, together with domain annotation. **(C)** The ratio of nonsynonymous to synonymous substitutions (dN/dS value, ω) of Hrr25p, by clade and domain. Domains are as in (C). Gene tree shown in **Supplementary Figure 1. (D)** Evolutionary distance among Hrr25p orthologs. (Left) Schematic of the gene tree. Branch lengths from the Pre-WGD node (shown in red) were computed using the tree based on non-synonymous changes (dN), with the tree structure shown in **Supplementary Figure 1**. (Right) Branch lengths from the Pre-WGD node, by maintenance status of *HRR25*. Mean ± sd is shown for each group. The difference in group means is compared with a two-sided Wilcoxon test. **(E)** Scatter plot representation of dN tree branch lengths for post-WGD species maintaining two ohnologs of *HRR25*. Grey dashed line represents the mean of post-WGD orthologs which reverted to single copy state. Black dashed line shows x=y. **(F)** Asymmetrical evolution of ohnologs by species. Rate asymmetry *R’* for each post-WGD species maintaining two copies of *HRR25*. *R’* values were computed as the ratio of values represented in (D) by dividing the value of the ohnolog with longer branch length with the ohnolog with the shorter branch length as in (Byrne and Wolfe 2007). Gray dashed line represents *R’* = 1, where evolutionary rates of both ohnologs are equivalent.

Although it is challenging to determine the exact mechanisms that led to the retention of a specific gene pair, each of these models make specific predictions regarding duplication, most of which have been tested in the budding yeast. The ancestor of some clades of the yeast family *Saccharomycetaceae* underwent whole genome duplication (WGD) that has impacted their cellular network architecture and functions. About 15% of ohnologs were maintained as pairs in the lineage leading to *Saccharomyces cerevisiae*, the other 85% have returned to single copy (Byrne and Wolfe 2005). The systematic study of the remaining ohnolog pairs revealed patterns of retention that are consistent with the various models outlined above. We will not review this rich literature here, but we will illustrate with a few cases.

A classical paper (van Hoof 2005) for instance tested the DDC model using complementation tests. In this model, duplicates are maintained by partitioning the ancestral function. The author showed that for at least three pairs of genes, the diverged functions of ohnologs could be complemented with an orthologous gene from a species that diverged prior to the whole genome duplication (WGD). One of the reasons why WGDs are powerful model systems for the study of gene duplication is that speciation events that occur after the WGD event lead to opportunities for repeated, partially independent gene pair maintenance in these species. For instance, the comparison of *S. cerevisiae* and *Kluyveromyces polysporus* (also known as *Vanderwaltozyma polyspora*) (Scannell et al. 2007) showed that genes associated with particular molecular functions were more or less likely to be retained after the WGD. One specific example was that both species showed an underrepresentation of RNA metabolism genes among the preserved ohnologs, but an over representation of protein kinases. In other studies, it was shown that data on the evolution of gene expression in various species support the model of dosage subfunctionalization, where duplicated genes show decreased expression levels, partitioning the dosage requirements between the two copies (Qian et al. 2010; Gout and Lynch 2015). They show that maintaining constant summed expression could result in neutral evolution, which gradually creates an imbalance of expression between ohnologs, leading to the nonfunctionalization of one copy.

Here, we examined in particular ohnologs that were maintained across many species, not focusing on their maintenance in *S. cerevisiae* alone. We searched for cases so that we could compare, for a given gene, multiple pairs of ohnologs that may be undergoing different fates in different species. We found one particular case, *HRR25*, that we examined in detail and combined multiple approaches to dissect their potential fate. The casein kinase (CK) 1δ/ε (gene *HRR25* in *S. cerevisiae*) is present in many eukaryotic proteomes where it phosphorylates serine or threonine residues on its numerous substrates (Tanaka et al. 2014). This kinase plays a role in a variety of functions including endocytosis (Peng, Grassart, et al. 2015), DNA damage repair (Hoekstra et al. 1991), ribosome biogenesis (Ghalei et al. 2015), meiosis (Ye et al. 2016; Zhang et al. 2018), autophagy (Pfaffenwimmer et al. 2014), and membrane trafficking (Wang et al. 2015). The deletion of this gene is reported to cause inviability in the laboratory strain of *S. cerevisiae* (Giaever et al. 2002), and mutations in the P-body localization signal sequence or heterozygous deletion of the entire gene result in the failure of spore formation (Zhang et al. 2018).

The maintenance of both *HRR25* duplicate copies and its large number of functions make it a good model to examine its pattern of molecular evolution following the yeast WGD. We examine whether the ohnologs have most likely been maintained by subfunctionalization or neofunctionalization, or if they may be slowly undergoing nonfunctionalization. To test the predictions of these various models, we examined the rate of evolution of the ohnologs and we performed functional complementation assays in *S. cerevisiae*, using the essentiality of the *S. cerevisiae* singleton as a phenotype. We did so by measuring the effects of their expression on cell viability while conditionally downregulating the endogenous *HRR25*. We investigated the molecular mechanisms for the observed viability by performing proteome wide protein-protein interaction (PPI) assays. The orthologs with similar PPI profiles to *S. cerevisiae* Hrr25p matched with that of the orthologs capable of complementation. Most of the ohnolog pairs exhibited one ohnolog that cannot complement and also drastically lost its PPIs, suggesting nonfunctionalization. Localization analysis also showed that the orthologs incapable of functional complementation show distinct patterns of subcellular localization, similar to that of a catalytic dead mutant reported in previous studies. We validated this finding in *N. castellii*, showing that one ohnolog of this species has become non-essential. Taken together, we show that ohnologs of *HRR25*, a kinase under strong negative selection when present in singleton, are undergoing asymmetrical evolution and have functional features that are consistent with their nonfunctionalization.

## Results and Discussion

### *HRR25* maintenance in post-WGD *Saccharomycetaceae* species are asymmetrical in evolutionary rates

Previous efforts have annotated the genomes of non and post-WGD species, and identified cases of gene maintenance and losses (Byrne and Wolfe 2005). Using this resource, we performed a preliminary analysis to search for cases of essential genes in *S. cerevisiae*, which reverted to the single copy state in this model species, but having both duplicated copies maintained in other post-WGD species. We found that *HRR25*, a gene coding for the kinase CK1δ/ε, is a rare case where both duplicated copies are maintained in 7 species post-WGD, which is much more than for other *S. cerevisiae* genes (Mean = 0.31, Z-score = 7.3, **Figure 1B**). Multiple sequence alignment of orthologs and subsequent residue similarity analysis against the ortholog of non-WGD species *T. delbrueckii* showed that species which maintained both *HRR25* ohnologs have lower sequence identity compared to non-WGD or single reverted state species (**Figure 1C**). This suggests that in species where both ohnologs are maintained, this kinase has been evolving faster.

We therefore examined the molecular evolution of orthologs by constructing a gene tree from protein sequences (**Supplementary Figure 1**). The ratio of non-synonymous to synonymous substitutions (dN/dS value, ω) was used to estimate the strength of selection acting on these genes. The ω value for *HRR25*, based on a homogeneous model, is 0.035, suggesting that it is under strong negative/purifying selection. We then asked if gene duplication relaxed the degree of purifying selection for *HRR25* orthologs. We examined this using a branch model to estimate an overall ω in post-WGD orthologs. We found that ω increased for 2 clades which contains orthologs from post-WGD species maintaining both ohnologs (ω = 0.058 & 0.056), compared to clades with post-WGD species that reverted to single copy (ω = 0.013), non-WGD species (ω = 0.011), and the outgroup (ω = 0.014) (**Supplementary Figure 1**). This shows that purifying selection has been relaxed specifically for orthologs in species which maintained both ohnologs.

In *S. cerevisiae*, *HRR25* has three domains: the kinase domain, the central domain, and the tail region (Corbett and Harrison 2012). The disordered C-terminal tail region has a high proline and glutamine content, and undergoes autophosphorylation to form a pseudosubstrate inhibitor that affects the activity and specificity of the kinase (Cullati et al. 2023). The central domain, packed tightly against the kinase C-lobe, is known to interact with other proteins, which is critical for the proper localization of the protein (Ye et al. 2016). We asked whether selection differs between the three domains. Relaxation of selection occurred most strongly on the disordered tail region, followed by the central and kinase domains (**Figure 1D**). In spite of its overall higher conservation level, both of the two post-WGD clades of species that maintained ohnologs show an increase of ω value in the kinase domain. On the contrary, one post-WGD clade (annotated as post-WGD maintained 2 in **Figure 1** and **Supplementary Figure 1**) showed larger and smaller ω values for the central and tail domains, respectively, compared to other clades.

The difference in evolutionary rates among post-WGD ohnologs was further assessed using branch lengths from the pre-WGD node to all extant orthologs in the gene tree, which was constructed from nonsynonymous substitutions (hereafter referred to as dN tree). The mean of branch lengths for orthologs from species that maintained both ohnologs post-WGD showed significantly larger values compared to non-WGD and single copy reverted post-WGD orthologs (mean ± sd; Non-WGD: 0.019 ± 0006; Post-WGD single copy: 0.062 ± 0.009; Post-WGD duplicate maintained: 0.227 ± 0.125, p-value ≤4.00 ×10^−9^, two-sided Wilcoxon test) (**Figure 1E**). The branch lengths for ohnolog pairs for species maintaining both duplicates post-WGD show that dN is variable between species and also between ohnolog pairs maintained in the same species (**Figure 1F**). To further examine the asymmetry, we measured rate asymmetry (*R′*) for each pair of ohnologs by taking the ratio of their branch lengths from the dN tree (Byrne and Wolfe 2007). The rate asymmetry measures of *HRR25* ohnologs pairs are highly variable among species, with *N. castellii* having the most asymmetrical rate of evolution among all ohnologous pairs (**Figure 1G**). To make sure that the high rate of evolution did not result from a sequencing error in the reference genome, we verified the coding sequence of *HRR25* in the *N. castelli* type strain using Sanger sequencing, which showed that the sequence was correct.

Reconciliation of the *HRR25* gene tree was performed to depict the relationships between non-WGD and post-WGD orthologs (see methods for details). Genes are annotated with gene names or encoded regions (See **Supplementary Table 1** for full detail). The node annotated pre-WGD was used to compute evolution rates in analysis shown in Figure 1.

### Combinatorial complementation assay reveal no subfunctionalization of *HRR25* post-WGD

The analyses of protein sequences show that the rate of evolution for *HRR25* post-WGD has globally increased in species in which both ohnologs are maintained and, and in many cases is asymmetrical between ohnologs. This increase in rate of evolution is most likely not a consequence of adaptive evolution, but of relaxed purifying selection, with a 5-fold increase in dN/dS but that remained below 1 (**Supplementary Figure 1**). This degree of relaxed selection, not reaching neutral evolution, is a trend also seen genome-wide for genes under negative selection post-WGD (Kondrashov et al. 2002).

We used a functional complementation assay to test if preserved ohnologs could perform the essential functions the gene has in *S. cerevisiae*. Each ortholog was expressed in *S. cerevisiae* to test if it can rescue the growth defect caused by the conditional repression of the *HRR25* (Figure 2A & **B**). We synthesized all yeast orthologs (n= 20, except *S. cerevisiae*) and cloned them in a CEN plasmid (low-copy number) using the same promoter for all genes so we can control for expression. For completeness, we also included 4 orthologs from *H. sapiens* as reports have shown that the human ortholog CSNK1E (but not CSNK1D nor CSNK1A1) can complement the *S. cerevisiae HRR25* (Laurent et al. 2020). We confirmed that all orthologs were expressed in *S. cerevisiae* using Western blotting (**Supplementary Figure 2**).

**Figure 2.**
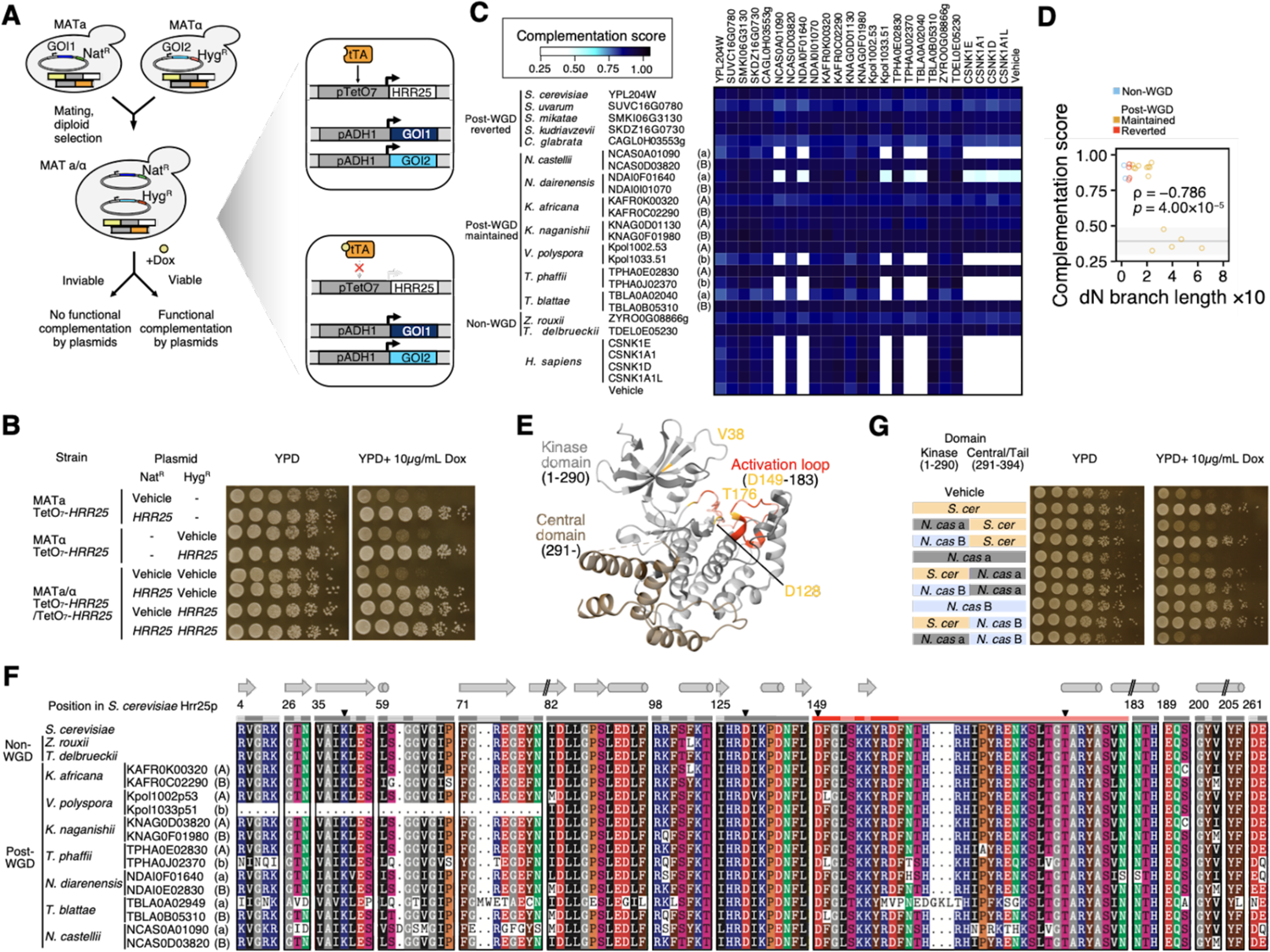
Most ohnolog pairs have one complementing and one non complementing gene, and the kinase domain is critical for this complementation. (A) The expression of the genomic *HRR25* in *S. cerevisiae* was put under the control of the TetO_7_ promoter. Gene expression under the TetO_7_ promoter requires tTA, which is inhibited in presence of doxycycline. Genes expressed on the CEN/ARS plasmids under ADH1 promoter are not affected by the presence of doxycycline. Functional complementation is assessed based on rescue of the inviable phenotype caused by repression of the endogenous *HRR25*, under presence of orthologs expressed on the CEN/ARS plasmid. (B) Functional complementation shown by spot dilution assays. Growth of diploid strains carrying CEN/ARS plasmids that constitutively express *HRR25* under the ADH1 promoter. Spot assay with vehicle and with the WT *HRR25* are shown as reference. In the absence of plasmid containing the WT *HRR25*, growth is severely diminished. Expressing *HRR25* from two plasmids does not diminish growth in a detectable manner. Spots were plated in 5-fold serial dilutions, starting from OD_600nm_ = 0.5. (C) Heatmap representation of complementation scores for combinations of orthologs. Complementation scores are relative growth, measured by growth curves of colonies on plates, under doxycycline divided by the YPD control (n=22 replicated colonies per genotype). Ohnologs are annotated either with A or B, corresponding to the genomic loci, based on the YGOB database. Non-complementing ohnologs are labeled with lower case (a or b). (D) Scatter plot of complementation scores and rates of evolution. Rates of evolution are dN values from the tree from the pre-WGD node (shown in Figure 1E). The gray line and shaded area indicate mean and 95% confidence interval for the Vehicle control, respectively. Value for *S. cerevisiae* was excluded from this analysis. Pearson correlation (*ρ* = −0.786, p-value = 4.00 × 10^−5^). (E) Crystal structure of the *S. cerevisiae* Hrr25p (PDB: 5CYZ, (Ye et al. 2016)). The kinase domain is colored gray. Known critical residues for catalytic activity are colored orange. The activation loop within the kinase domain is colored red. The central domain is colored brown. Darker shades within the kinase domain represent positions that are conserved in the *S.cerevisiae* kinases *HRR25*, *YCK1*, *YCK2*, and *YCK3* (Murakami et al. 1999). (F) Multiple sequence alignments of ohnologs showing the conserved residues in the kinase domain. Post-WGD species are ordered based on evolutionary asymmetry (*R’*) values. Protein positions for *S. cerevisiae* are shown on top. Positions which correspond to helix or sheet structures are annotated above positions. Critical residues for catalytic activity are annotated with black triangles. Conservation of residues within *S. cerevisiae* are shown under the protein positions, with the color code as in (E). (G) Spot dilution assay to test for the complementation of chimeric genes to assess the role of the kinase domain in complementing the *S. cerevisiae HRR25*. The kinase domain (positions corresponding to 1-290 in *S. cerevisiae HRR25*) were swapped between *S. cerevisiae* and *N. castellii* orthologs. *S.cer*: *S. cerevisiae HRR25*; *N. cas* (a): *N. castellii* NCAS0A01090 (a); *N. cas* (B): *N. castellii* NCAS0D03820 (B). Cells were spotted in 5-fold dilution series, starting at OD_600nm_ = 0.5.

If the orthologs underwent subfunctionalization of essential functions, growth rescue will be achieved only when expressing both orthologs within pairs since they are expected to have complementary functions. Furthermore, if subfunctionalization took place for the same functions in different post-WGD species, orthologs from two species could also have complementary functions and be capable of complementing the function of *HRR25* as a pair. For cases where ohnolog pairs have undergone asymmetrical evolution through relaxed selection (e.g. *N. castellii*, *T. blattae*, etc.), we expect the copy which evolved faster to fail to complement, if the rapid evolution is leading to nonfunctionalization. Cases where both ohnologs are evolving at similar rates (e.g. *K. africana*, *K. naganishii*, etc.) are both expected to show functional complementation, since they are more likely to maintain the ancestral functions. If orthologs have undergone neofunctionalization, only one of the ohnologs, which kept the ancestral function, will achieve complementation if functions were lost in the ohnolog that gained a new function. If the gain of functions were not accompanied with loss of other functions, both ohnologs should complement. In order to test all of these predictions, functional complementation was assayed in a combinatorial manner, testing two orthologs encoded on separate plasmids in the same cell. In order to easily associate genes to their ability to functionally complement, genes were annotated with the encoded loci (e.g. A and B) from the YGOB database (Byrne and Wolfe 2005) in upper or lower cases, to describe functional complementation and failure of complementation, respectively.

First, we found that all single copy orthologs, including orthologs from the species that did not undergo WGD (*Z. rouxii* and *T. delbrueckii*) fully complement the repression of *HRR25* in *S. cerevisiae*, just as the *S. cerevisiae* gene does. This confirms that the function that is now essential is *S. cerevisiae* was present in the common ancestor of the post-WGD species. We observed no pair dependent growth complementation, which was expected in the case of subfunctionalization where ohnologs within pairs would have complementary functions (Figure 2C). Instead, for species that maintained both ohnologs, at least one could, but most often not both, complement the function of the *S. cerevisiae HRR25*. The orthologs which failed to complement were ohnologs from *N. castellii*, *T. blattae*, *N. dairnensis*, *V. polyspora*, and *T. phaffii*. Such orthologs except for *V. polyspora* have all evolved faster than their counterpart ohnolog, and are likely evolving under relaxed selection. Species for which ohnologs were evolving at similar rates (*K. africana*, *K. naganishii*, and *V. polyspora*) showed complementation for both orthologs, except for *V. polyspora*. The non complementing *V. polyspora* ohnolog, Kpol1033p51 (b), has a frameshift mutation likely causing the mRNA to translate from an alternative start codon, resulting in a partial kinase domain.

We did not observe any complementation from any of the human orthologs. The failure of complementation for CNSK1E, which was previously reported to complement, can be attributed to the difference in experimental settings such as the promoter and the exact isoforms used for the assay. We therefore focused only on the yeast orthologs for the downstream analysis. Globally, we observed a negative correlation between evolutionary rate and complementation score (see methods), confirming further the association between the rate of evolution and the preservation of the ancestral functions (Figure 2D, Pearson correlation coefficient, *ρ* = −0.786, p-value = 4.00 × 10^−5^).

We asked if the failure of complementation can be attributed to the loss of catalytic activity of the kinase domain, since it is the protein domain with an enzymatic activity. We investigated critical residues whose mutations cause the kinase to become pseudokinases (Kung and Jura 2019). In *HRR25*, these residues correspond to the β3 lysine (V38) within the VAIK motif, the aspartate (D128) in the HRD motif, and the aspartate (D149) in the DFG motif. Additionally, in *HRR25*, the mutant T176I has been reported to create a kinase dead mutant (Murakami et al. 1999). A previous study in the mammalian homolog CK1δ (CSNK1D) has reported that similar mutations (K38M and T176I) alter the localization of the protein (Milne et al. 2001) (Figure 2E). The post-WGD orthologs did not show mutations in these motifs, but rather showed accumulation of mutations in the activation loop and substrate-binding regions, including positions that are conserved among *S. cerevisiae* kinases (*YCK1*, *YCK2*, *YCK3*, and *HRR25*) (Figure 2F). We therefore hypothesized that the evolution of the kinase domain could be sufficient to cause the incapability for functional complementation. We focused on *S. cerevisiae* and *N. castellii* orthologs, since *N. castellii* ohnologs have shown the most asymmetrical evolution among post-WGD species. We constructed chimeric *HRR25* genes by swapping the kinase domain between *S. cerevisiae* and *N. castellii* orthologs. Complementation assay of chimeric subunits showed no complementation for all constructs containing the kinase domain of the non-complementing *N. castellii* NCAS0A01090 (a), while the central and tail domains did not alter the functional complementation of *S. cerevisiae* or the functional *N. castellii* NCAS0D03820 (B) (Figure 2F). This illustrates that the complementation capability of the *N.castelli* ohnologs are a result of differences in kinase domain activity. Taken together, we observed results that are overall consistent with the nonfunctionalization of many *HRR25 ohnologs*, but not subfunctionalization nor neofunctionalization at the gene level.

### Proteome wide protein-protein interaction profiles reveal some gains but mostly loss of protein-protein interactions

We wanted to further examine the functionality of *HRR25* ohnologs. For this, we mapped protein-protein interactions (PPIs) as the patterns of PPI partners are a powerful indicator for the function of proteins (Marcotte et al. 1999; Mayer and Hieter 2000; Hishigaki et al. 2001).

We performed a proteome-wide DHFR-PCA screening using *HRR25* orthologs as bait and *S. cerevisiae* proteins as preys. We focused on all *S. cerevisiae* proteins that have been shown to have at least one PPI with this experimental approach (Tarassov et al. 2008; Rochette et al. 2015; Diss et al. 2017). Among the 1,172 *S. cerevisiae* proteins subjected for the screening, 1,085 proteins passed the quality check for interaction evaluation (see methods for details). We identified 115 proteins that interacted with at least one *HRR25* ortholog (Figure 3A). We performed gene ontology (GO) enrichment analysis on the detected interaction partners. Significant enrichment was observed for GO terms with descriptions of cytoskeleton, cell tip, endocytotic patch, vesicle, and RNA (P-body, Polysome, and U6 snRNP) (Figure 3B), which agrees with previous reports for Hrr25p function (Peng, Moritz, et al. 2015; Peng, Grassart, et al. 2015; Wang et al. 2015; Zhang et al. 2016).

**Figure 3.**
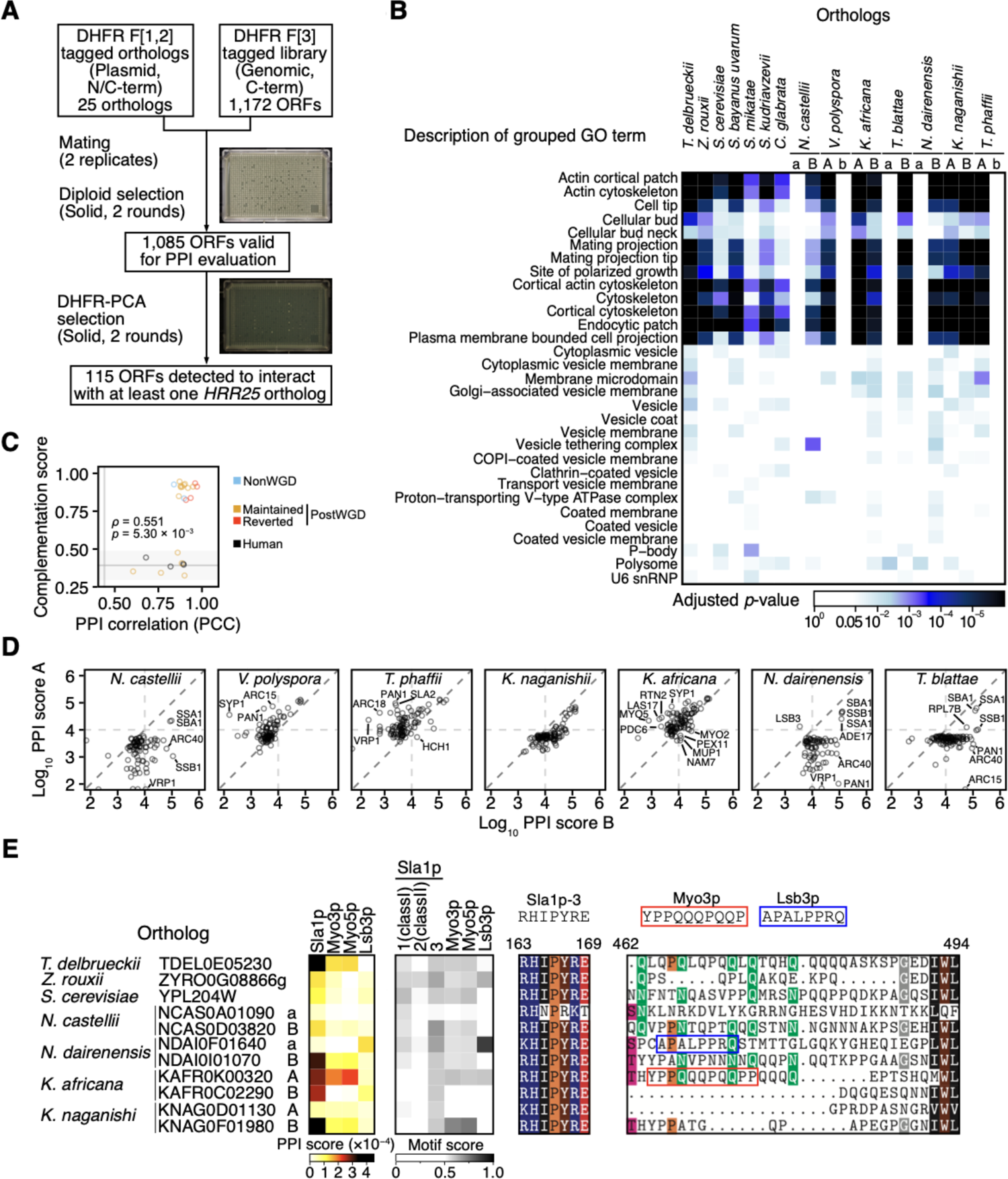
Protein interaction profiles of Hrr25 orthologs with *S. cerevisiae* proteins. (A) Experimental workflow for proteome-wide PPI screening. *HRR25* orthologs were cloned to CEN/ARS plasmids, and subjected for screening against the proteome-wide genomically tagged DHFR F[3] collection for DHFR-PCA screening. Typical plate images of endpoints are shown for diploid selection and PPI selection. (B) Heatmap representation of GO enrichment of detected PPIs for each ortholog. GO terms with the same descriptions were grouped. Lowest values for each GO term description were adopted for visualization. See **Supplementary Table 7** for detailed description and raw data. (C) Correlation between PPI profile similarity against *S. cerevisiae* and functional complementation scores. PPI profile similarity values are Pearson correlation coefficients of a given ortholog versus the *S. cerevisiae* ortholog using the PPI score of 115 PPI partners. The gray line and shaded area indicate mean and 95% confidence interval for the Vehicle, respectively. Values for *S. cerevisiae* were excluded from this analysis. Pearson correlation (*ρ* = 0.551, p-value = 5.03 × 10^−3^). (D) Scatter plot comparison of PPI scores for maintained ohnologs pairs in post-WGD species. Horizontal and vertical dashed lines represent thresholds to call positive interactions. The diagonal line represents equal PPI scores for both ohnologs. (E) PPI profiles for selected orthologs on SH3-encoding protein interaction partners which showed loss or gain of PPIs. Left: Heatmap representation of PPI scores. Middle: Heatmap representation of motif scores for each binding motif within orthologs. Right: MSA of binding regions for Sla1p, Myo3p, and Lsb3p. Binding motifs within the region are indicated above the MSA. Protein positions for *S. cerevisiae* (total length of 494) are shown above the MSA.

We observed major differences in GO enrichment between ohnolog pairs, due to loss of individual PPIs for the non-complementing and rapidly evolving ohnologs, namely for *N. castellii* NCAS0A01090 (a), *V. polyspora* Kpol1033p51 (b), *T. blattae* TBLA0A02040 (a), *N.dairenensis* NDAI0F01640 (a), and *T. phaffii* TPHA0J02370 (b). This contrasts with the matching complementing ohnologs, which showed similar GO enrichment profiles as the non-WGD and single copy post-WGD orthologs. The similarity of PPI profiles compared with the *S. cerevisiae* Hrr25p showed a positive correlation with the functional complementation (Pearson correlation coefficient, *ρ* = 0.551, *p*-value = 5.30 × 10^-3^), which indicates that the PPI differences, mainly the losses, mirror the ability to complement (Figure 3C). In most cases, ohnologs maintained in post-WGD species showed a strong asymmetry of PPIs, where one copy of the post-WGD ohnolog pair has lost most of its PPIs (Figure 3D). In *N. castellii*, most PPIs were maintained for the ohnolog NCAS0D03820 (B), whereas the other ohnolog NCAS0A01090 (a) only showed interactions with Ssa1p and Sba1p. These two proteins involved in protein folding are abundant proteins that are flagged as non-specific interactors in DHFR-PCA screens and that are used as abundance controls (Tarassov et al. 2008). Detecting these interactions confirmed the protein level expression of ohnologs, even for copies which were incapable of functional complementation. These analyses strongly support the previous results that many of the faster evolving ohnologs are on their way to nonfunctionalization as they lost many functions related to their interaction with other proteins.

One exception that could potentially show divergence of function is ohnologs maintained in *K. africana*, which showed ohnolog specific PPIs. Such PPIs were in some cases not observed for non-WGD orthologs or the *S. cerevisiae* ortholog. These could be cases of interactions that evolved in this species since the WGD event. Among all ohnolog-specific PPIs, we focus on interactions that have similar molecular functions. We observed two such cases, Pex11p/Pex19p/Pex30p and Myo2p/Myo3p/Myo5p (Figure 3D). In the case of Pex11p/Pex19p/Pex30p interactions, we observed both ohnologs interacting with Pex19 as in non-WGD orthologs, whereas ohnologs KAFR0C02290 (B) and KAFR0K00320 (A) separately gained interactions with Pex11p and Pex30p, respectively. Pex11p, Pex19p, and Pex30p are proteins involved in the peroxisome biogenesis. While Pex19p plays a role in the docking of the receptor–cargo protein complex on the peroxisome membranes, Pex30p and Pex11p are involved in distinct timing and processes of the peroxisome biogenesis pathway. More specifically, Pex30p localizes to the ER, enriched at the site of pre-peroxisome vesicle formation, and regulates the tubular structure. Pex11p is responsible for membrane elongation of pre-existing peroxisomes (Deori and Nagotu 2022). Since non-WGD orthologs show interactions with some of these proteins, the detection of specific protein-protein interactions for KAFR0C02290 (B) and KAFR0K00320 (A) with them may indicate slight changes of function.

In the case of Myo2p/Myo3p/Myo5p interactions, we observe that the ohnolog KAFR0K00320 (A) kept the interaction with Myo3p/Myo5p as in non-WGD orthologs, while the other ohnolog KAFR0C02290 (B) has lost the interaction with Myo3p/Myo5p and gained interaction with Myo2p, which was not detected for any of the non-WGD orthologs. Myo2p, Myo3p, and Myo5p are myosin motor proteins, and they are involved in actin-based cargo transport and endo/exo-cytosis in actin cortical patches, respectively (Tanaka and Matsui 2001). Myo3p and Myo5p contain a Src homology 3 (SH3) domain, which contributes to PPIs by recognition of binding motifs (Tonikian et al. 2009; Dionne et al. 2021). The SH3 domains in Myo3p and Myo5p are required for targeting to sites of actin polarization (Anderson et al. 1998). Interestingly, Myo2p is also an ohnolog (with Myo4p) so this change in PPIs could have evolved since the WGD. When focusing on other PPI partners containing SH3-domains, we found that Lsb3p and Sla1p gained and lost PPIs in some orthologs, respectively.

PPIs with SH3-domains are achieved through binding motifs, leading us to examine the sequence difference in orthologs contribute to the gain and loss of PPIs against SH3-domain containing proteins. We examined the presence of SH3 binding motifs in *HRR25* orthologs for the SH3-domain containing proteins of which we observed at least one interaction in this study. To identify the SH3 binding motifs, we used position weight matrices (PWMs) constructed from quantitative motif binding assays (Tonikian et al. 2009). We examined the occurrence of PWM matches within each ortholog (MSS; matrix similarity score), and used the highest value as the motif score (see methods for detail). We found that the *K. africana* ohnolog, KAFR0C02290 (B), which lost interactions with Sla1p, Myo3p, and Myo5p has acquired mutations contributing to the decrease of motif score for the corresponding binding partners. In contrast, the *N. dairenensis* ohnolog NDAI0F01640 (a), which gained interaction against Lsb3p, has accumulated mutations which increased the motif score for the Lsb3p binding motif (Figure 3E).

Taken together, while neofunctionalization and/or subfunctionalization at the GO term level were not observed, we observed instances of gain of specific PPIs in two separate biological processes in the *HRR25* orthologs post-WGD. In both of these cases, the gained interaction partners were both functionally related to an interactor of non-WGD orthologs. These cases could represent cases of neofunctionalization in the classical sense (Ohno 1970). The potential adaptive significance of these changes remains to be investigated.

### Non-complementing ohnologs have abnormal subcellular localization

Hrr25p is known to require both kinase activity and PPIs for its functionality. For example, the central domain of Hrr25p interacts with Mam1p, localizing Hrr25p to the meiosis I centromeres (Ye et al. 2016). In this case, both kinase activity and Mam1p binding are essential for monopolar attachment of Hrr25p, shown from abnormal chromosome segregation when mutations in the kinase domain (H25R and E34K) were introduced (Petronczki et al. 2006). Mutations that disrupt the kinase function also cause abnormality in localization for the human CK1δ (CSNK1D) (Milne et al. 2001). Another example is the localization of Hrr25p to cytoplasmic RNP granules (P-bodies) under stress conditions and during meiosis. The localization of Hrr25p to P-bodies protects this enzyme from the degradation machinery during these periods of stress (Zhang et al. 2016). The P-body localization signal (PLS) in the central domain mediates the interaction between Dcp2p and allows Hrr25p to localize to the P-body. This localization of the P-body foci is also seen for the human CK1δ (CSNK1D) in HeLa cells (Zhang et al. 2016). While kinase activity is required for P-body localization of Hrr25p, the mutations that cause deficiency in monopolar attachment (H25R and E34K) do not affect localization to P-body foci or the sporulation efficiency (Zhang et al. 2018). Collectively, these observations indicate that the diverse functions of Hrr25p are achieved from independent functionalities represented through kinase activity and/or PPIs.

We examined the localization of each ortholog using an mEGFP reporter (**Supplementary Figure 3**). The *S. cerevisiae* Hrr25p showed localization similar to previous observations, localizing to the budneck, cytoplasm, nucleus, and the bud (Ghaemmaghami et al. 2003). The orthologs from non-WGD species, *Z. rouxii* and *T. delbrueckii*, showed similar localization patterns to the *S. cerevisiae* Hrr25p, again showing that this is likely the ancestral localization. The localization of complementing ohnologs also showed similar patterns to that of the *S. cerevisiae* Hrr25p. On the other hand, consistent with the complementation assays, non-complementing ohnologs did not show localization to the budneck, but instead showed localization constituting foci in cytoplasm, while the *T. blattae* ortholog TBLA0A02040 (a) showed exclusive localization to the nucleus, similar to the report for the catalytic dead human homolog (Milne et al. 2001). The localization of orthologs agrees with our observations on functional complementation and PPIs.

The illustration of the plasmid design for mEGFP fusion of *HRR25* orthologs is shown on the top left corner. Cells under vegetative growth were imaged using confocal microscopy (see methods for details). For post-WGD orthologs the genes are annotated using functional complementation results, using upper and lower case for functional and non-functional orthologs, respectively. Scale bar represents 10 µm.

### One ohnolog appears on its way to nonfunctionalization in *N. castellii*

We have shown that many *HRR25* ohnologs in post-WGD species underwent asymmetrical evolution and lost the ability to complement the essential function of this kinase. These results suggest that one gene among the ohnolog pairs are on their way to nonfunctionalizaiton. One striking case is the *N. castellii* ohnolog NCAS0A01090 (a) compared to its ohnolog NCAS0D03820 (B). Sequence level analysis has shown that this ohnolog has evolved faster than the other ohnolog, under relaxed selection. This ortholog has failed to complement the loss of *HRR25* in *S. cerevisiae*, lost most of the PPIs we observed, and showed abnormal subcellular localization. Changes in the kinase domain of NCAS0A01090 (a) specifically appear to be causing these losses of function.

These observations led us to hypothesized that *HRR25* in post-WGD species, at least in *N. castelli*, are evolving under the dosage subtunfcitonalization model (Gout and Lynch 2015), where eventual decrease in gene expression promotes nonfunctionalization of one ohnolog by relaxing selection on its function. We first examined the expression level of *HRR25* orthologs across non-WGD and post-WGD species. We compared expression data from *S. cerevisiae* (post-WGD, single copy reverted), *N. castellii* (post-WGD), and *Kluyveromyces lactis* (non-WGD), and normalized the data based on common 1:1 orthologs to be able to directly compare transcript levels among species, as done in (Qian et al. 2010). Based on available gene expression data, we observed that *HRR25* expression levels in *S. cerevisiae* and *K. lactis* are slightly above the mean of 1:1 orthologs (Log_2_ fold-change of 0.568 and 0.140, respectively). In *N. castellii*, the complementing ohnolog NCAS0D03820 (B) has a similar level of expression to the mean 1:1 orthologs (Log_2_ fold-change = −0.035) (Figure 4A). Contrastingly, the ohnolog NCAS0D03820 (a) has a lower expression level, with a Log_2_ fold-change of −2.154 relative to the mean of 1:1 orthologs. We examined how this difference in expression compares with the divergence of expression of other ohnolog pairs and found that the expression difference between NCAS0A01090 (a) and NCAS0D03820 (B), a 5-fold difference in expression, is at the 70^th^ percentile in expression asymmetry among all WGD duplicate pairs in this species (Figure 4B **& C**). Interestingly, a >5-fold difference in expression was previously shown to cause an asymmetry of mutational effects, where new nonsynonymous mutations and frameshift-causing indels are significantly more deleterious in the highly expressed copy compared with their paralogs with lower expression (Johri et al. 2022), which is consistent with our observation of nonfunctionalization.

**Figure 4.**
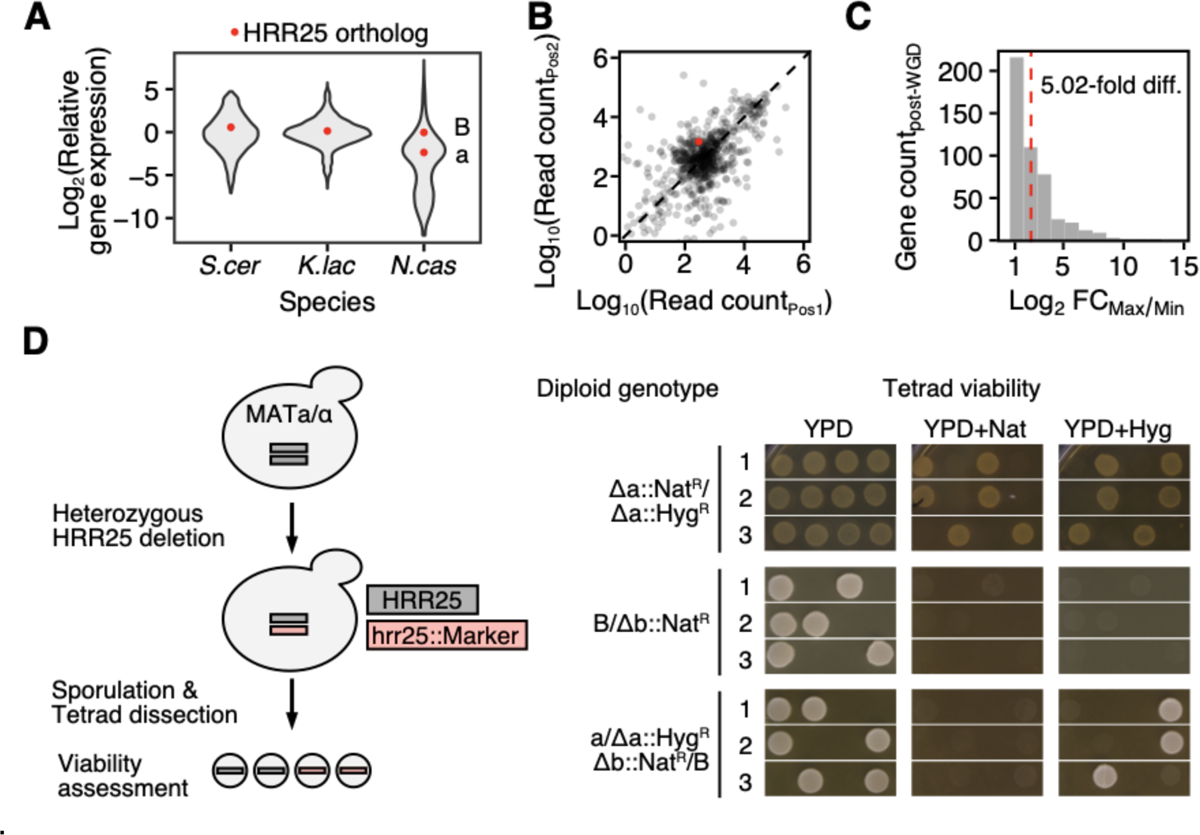
Validation of the loss of essential function for *N. castellii* NCAS0D03820. (A) *HRR25* gene expression in *S. cerevisiae*, *K. lactis*, and *N. castellii*. Violin plots represent the distribution of gene expression levels in each species. Expression values are normalized based on mean expression levels of 1:1 orthologs present in all 3 species. Red points indicate the *HRR25* orthologs in each species. *N. castellii* ohnologs are annotated as; a (NCAS0A01090) and B (NCAS0D03820). (B) Scatter plot representation of gene expression for post-WGD ohnolog pairs in *N. castellii*. Each point represents one ohnolog pair, with raw read counts retrieved from GSE90041. The diagonal line represents equal expression between ohnolog pairs. Red point denotes the ohnolog pair which is orthologous to *HRR25*. The axes (Pos1 and Pos2) are interchangeable for each point and do not reflect any meaning for the encoded loci. (C) Distribution of gene expression imbalance for post-WGD ohnolog pairs in *N. castellii*. Log_2_ fold-change (FC) was computed for each WGD ohnolog pair by taking the ratio of read counts between the higher (Max) and lower (Min) ohnolog. Red dashed line represents the value for the *HRR25* orthologs, with a fold change value of 5.02. (D) Tetrad dissection analysis in *N. castellii* for evaluating the essentiality of each *HRR25* ohnolog. (Left) Illustration of the experimental procedure. Diploid *N. castellii* strains were used to construct strains carrying heterozygous deletion of *HRR25* ohnologs. Strains were then sporulated and tetrads were dissected for assaying the viability of reach genotype using selection markers. (Right) Spot assay for tetrad viability. Three diploid strains were subjected for this analysis. Each row represents a different tetrad. Three of the typical tetrads are shown. *HRR25* loci are annotated as (a) NCAS0A01090 and (B) NCAS0D03820.

To validate our finding that NCAS0A01090 (a) has lost the ancestral essential function and is likely undergoing nonfunctionalization, we tested the essentiality of these two *N. castelli* ohnologs *in vivo*. We knocked out the orthologs in *N. castellii*, and performed tetrad dissections to assess spore viability. As a result, tetrads with the NCAS0A01090 (a) knockout but not NCAS0D03820 (B) knockout were viable, indicating that NCAS0A01090 (a) has become dispensable in *N. castellii* (Figure 4D). The imbalance of essentiality demonstrates that the ancestral essential functions of *HRR25* are maintained only in NCAS0D03820 (B), but have been lost for NCAS0A01090 (a).

## Conclusion

In this study, we investigated the fate of *HRR25* using orthologs from 12 species descending from the same WGD event. We asked which of the evolutionary scenarios (subfunctionalization, neofunctionalization, or nonfunctionalization) could explain the retention of *HRR25* duplicates in many of these species. Sequence level analysis showed that *HRR25* in species which maintain both ohnologs is under more relaxed selection, compared to orthologs of non-WGD or single copy reverted post-WGD species. Functional complementation assays showed that when ohnologs are evolving asymmetrically, the faster evolving ohnolog fails to complement the function of the *S. cerevisiae HRR25*. A proteome-wide PPI screen against *S. cerevisiae* proteins showed similar patterns at the molecular level, where orthologs evolving faster than other orthologs have lost most of their PPIs. We validated our findings in *N. castellii*, and showed that the faster evolving ohnolog has become dispensable in this species. The fact that this dispensable copy has much lower expression *in vivo* supports its potential decreased functionality. Taken together, our results indicate that many *HRR25* orthologs could be nonfunctionalizing, likely following the dosage subtunfcitonalization model (Gout and Lynch 2015), whereby asymmetry in expression leads to relaxed selection on one ohnolog and then to its slow degeneration at the protein level. This model does not apply to all WGD species. For instance, in some species where both ohnologs of *HRR25* appear more conserved, we discovered potentially recently gained PPIs, which may suggest neofunctionalization of some ohnologs that is supported by the coincidental evolution of SH3-domain binding motifs. Such gain of PPIs may represent early states of neofunctionalization in the classical sense (Ohno 1970). The contribution of these gained PPIs on any biological functions and/or adaptive significance remains to be explored.

We recognize the limitations of our work. Namely, because all other species are not amenable to similar assays, complementation and PPI assays were performed in *S. cerevisiae*. The complementation assays could only reveal results related to the essential functions of the *S. cerevisiae* ortholog and not other functions that could have evolved in other species. This is also the case for PPIs. In addition, the co-evolution of the PPI partners within species could lead to losses of interactions in *S. cerevisiae* even if PPIs are maintained within the other species. However, this would be unlikely to occur at the scale of the entire interactome. The support to our conclusions comes from the combination of various observations made based on the rates of evolution and the assays we performed. *HRR25* represents a rare case of maintenance of ohnolog pairs in many post-WGD species and as such, it could be an exceptional representative of pairs that are destined to return to single-copy but that have not completed the process yet. One potential question that our results lead to is, given the ease with which mutations can lead to complete loss of function of genes, for instance through indels and stop codons, why are *HRR25* ohnologs sequences still largely intact? One potential answer is that the dosage subfunctionalization model leads to a slow decay of gene functions as changes in expression and coding sequences slowly reinforce each other. By reducing the expression level of one ohnolog, early regulatory mutations could have reduced the strength of selection acting on the coding sequence, leading to the accumulation of degenerative mutations that would have been deleterious in the ancestral copy or in the other ohnologs. A less functional protein would then contribute less to function, potentially lessening selective pressures on expression level on this ohnolog. This feedback cycle, which eventually will lead to gene loss being effectively neutral (Gout and Lynch 2015), could be very slow and *HRR25* could be one of the last of these pairs of genes undergoing this process.

## Material and Methods

### Ortholog sequence retrieval

We first extracted protein sequences of *HRR25* for species in the *Saccharomycetaceae* from the NCBI RefSeq database (https://www.ncbi.nlm.nih.gov/refseq, last accessed on Sep 18^th^, 2023). We used the phylogenetic tree from (Li et al. 2021) as the species reference tree. We trimmed this species tree based on the species of which the *HRR25* protein sequences were retrieved, using the tree_subset function of the R package treeio (Wang et al. 2020). For additional species which were not included in our first criteria but present in the trimmed tree, genome sequences were retrieved from the NCBI database and (Shen et al. 2018). *HRR25* orthologs were identified by aligning orthologs from the first criteria to the genomic sequences using BLAST (version 2.6.0+) (Altschul et al. 1990), followed by precise extraction of *HRR25* loci based on a global alignment using MAFFT (version 7.392) with the L-INS-i method (Katoh and Standley 2013). We manually checked the resulting dataset, and observed that in some species not all orthologs had been retrieved due to low genomic coverage, as seen from orthologs partially aligning to the edge of contigs. We therefore excluded *Kazachstania* species with only one *HRR25* ortholog identified for further analysis, eliminating single copy reversion annotations which are not reflecting the actual events. The annotated sequences of *N. castellii* orthologs in the type strain CBS 4309 (Wolfe et al. 2015) were validated by Sanger sequencing. The details of genes used in this analysis are shown in **Supplementary Table 1**.

### Multiple Sequence Alignment analysis

The global multiple sequence alignment of *HRR25* orthologs was performed using MAFFT (version 7.392) with the L-INS-i method (Katoh and Standley 2013). A custom script was used to evaluate the residue similarity against the non-WGD ortholog of *T. delbrueckii.* For each of the orthologs, we obtained the similarity score for each residue using the BLOSUM62 similarity matrix (Henikoff and Henikoff 1992). Indels were assigned a score of −10 for this analysis. We computed the normalized relative conservation score of each residue by dividing the similarity score with that of an identical match against *T. delbrueckii*, and multiplied this value by 100.

### dN/dS analysis

We constructed a reconciled gene tree using the species tree from (Li et al. 2021), and replaced the WGD clade using the gene tree of post-WGD orthologs. The gene tree for post-WGD duplicates were created following the tutorial paper by (Álvarez-Carretero et al. 2023), using codon based alignment using TranslatorX (Abascal et al. 2010), and RAxML-NG (Kozlov et al. 2019). For the dN/dS analysis, we eliminated aligned positions which were not present in more than 90% of orthologs. The dN/dS values were computed using the codeml program in the PAML package (version 4.10.6, Last accessed on Sep 18^th^ 2023 at https://github.com/abacus-gene/paml) (Yang 2007). The homogenous and branch models were used. For domain specific computation of dN/dS, we separately executed the codeml program using the branch model on each of the three protein domains. Domains for each ortholog were inferred from alignment to the *S. cerevisiae HRR25*, where they are defined as kinase domain (1-290), central domain (291-394), and the disordered tail region (395-). Evolutionary rates were referred to the branch lengths of the dN tree output from the codeml program. Distances between nodes were calculated using the get_all_pairwise_distances function of the R package castor (version 1.7.11) (Louca and Doebeli 2018).

### List of strains and plasmids used in this study

The detailed list of biological resources used in this study are listed in **Supplementary Table 2**.

### Oligonucleotides and gene fragments used in this study

Oligonucleotides and gene fragments used in this study are listed in **Supplementary Table 3**.

### Yeast medium

YPD medium was used for general yeast cultures and complementation assays. We used buffered SC medium at pH 6.0 (amino acid dropout mix, 2.0% glucose, 0.1% monosodium glutamate, 1% succinic acid, and 0.6% NaOH) for yeast cultures for Western blotting and microscopy experiments. Antibiotics were supplemented in final concentrations of G418: 200 µg/mL, CloNAT (NAT): 100 µg/mL, Hygromycin B (HPH): 250 µg/mL, and doxycycline (DOX): 10 µg/mL, unless otherwise stated.

### Strains construction for combinatorial functional complementation assay

The background strain R1158 was used for constructing the doxycycline inducible repressor strain (Horizon discovery, YSC1210) (Hughes et al. 2000). In order to enable efficient combinatorial screenings via mating, we constructed a BY4742 derived MATα strain for constructing doxycycline repressor strains. We achieved this by amplifying the URA3::CMV-tTA cassette from R1158 using primers CLO1-6 and CLOP167-D11. Amplification was performed using Q5 High-Fidelity DNA Polymerase (New England Biolabs, M0491L). The PCR reaction, in a final volume of 25 µL, consisted of 1X Q5 Reaction buffer, 200 µM of dNTPs, 500 nM of each forward and reverse primer, 0.5 unit of Q5 polymerase, and 200 ng of template genomic DNA. Thermal cycling conditions were (i) 98 °C for 30 s; (ii) 35 cycles of 98 °C for 10 s, 64 °C for 10 s, and 72 °C for 4 min; (iii) 72 °C for 10 min; and (iv) 12 °C forever. We integrated this cassette to the ura3 locus of BY4742 by homologous recombination using standard lithium acetate transformation, and selected the successful transformants on SC-Ura plates. The resulting strain yPE014 was subjected to genotyping PCR for confirmation using the same conditions as above.

### Plasmid construction for tet-off repressor strain construction

For efficient construction of TetO_7_ promoter tagged strains, we created a plasmid, pKB33, having the KanMX-TetO_7_ cassette (Hughes et al. 2000) in the cloning vector pUC19. We amplified the genomic region encoding the KanMX-TetO_7_ from R1158 derived yTHC4703 (Horizon Discovery, YSC1180) using primers CLOP258-A6 and CLOP258-B6, and inserted to the pUC19 digested with restriction enzymes SacI-HF (New England Biolabs, R3156) and XbaI (New England Biolabs, R0145S) by *in vitro* DNA assembly (Gibson et al. 2009). This product was transformed into chemically competent *E. coli* strain MC1061 and selected on 2YT+Amp plates. The construct was validated using genotyping PCR followed by Sanger sequencing of the inserted region.

### Tet-Off repressor strain construction

The KanMX-TetO_7_ cassette on the plasmid pKB33 was PCR amplified with primers CLOP280-C5 and CLOP280-D5 having homology to replace the promoter region of *HRR25*. Amplification was performed using Kapa HiFi Hotstart Polymerase (Roche, 07958897001). The PCR reaction, in a final volume of 25 µL, consisted of 1X reaction buffer, 300 µM of dNTPs, 300 nM of each forward and reverse primer, 0.5 unit of polymerase, and 1ng of template DNA. Thermal cycling conditions were (i) 95 °C for 5 min; (ii) 35 cycles of 98 °C for 20 s, 60 °C for 15 s, and 72 °C for 1:45 min; (iii) 72 °C for 3:30 min; and (iv) 12 °C forever. The resulting PCR product was used for genomic integration of yeast strains R1158 and yPE014, and plated on YPD+G418 plates for successful transformants. Insertion of the KanMX-TetO_7_ cassette was confirmed by genotyping PCR using primers CLOP280-A5 and CLOP280-B5 using thermal cycler conditions as described above. We named the R1158 derived MATa strain GS007, and the yPE014 derived MATα strain GS009.

### Gene synthesis and ENTRY clone preparation for Gateway cloning

We amplified the *S. cerevisiae HRR25* from genomic DNA of BY4741 using primers CLOP280-E5 and CLOP280-F5 to add the attB sites for Gateway BP reactions. Amplification was performed using Kapa HiFi Hotstart Polymerase (Roche, 07958897001). The PCR reaction, in a final volume of 25 µL, consisted of 1X reaction buffer, 300 µM of dNTPs, 300 nM of each forward and reverse primer, 0.5 unit of polymerase, and 4 ng of template DNA. Thermal cycling conditions were (i) 95 °C for 5 min; (ii) 30 cycles of 98 °C for 20 s, 60 °C for 15 s, and 72 °C for 1 min; (iii) 72 °C for 3:30 min; and (iv) 12 °C forever. For orthologous genes, we focused on species present in the YGOB database (Byrne and Wolfe 2005), with available RefSeq protein records. Protein sequences of orthologs were retrieved from the RefSeq protein files, and DNA fragments were synthesized (TWIST biosciences). All DNA fragments were codon optimized for expression in *S. cerevisiae*, and attB sequences were added for Gateway BP cloning. For genes which exceeded the synthesis limit (TBLA0A02040 and NDAI0F01640), we added the attB sites via PCR. PCR reactions were performed using the same conditions as described above. Primer information are listed in **Supplementary Table 3**. We individually subcloned the *HRR25* orthologs to pDONR201 by Gateway BP cloning (Invitrogen, 11789), following the manufacturer’s instructions. This product was transformed to chemically competent *E. coli* strain DH5α and selected on LB+Kan plates. The constructs were validated by Sanger sequencing. The human orthologs were retrieved from the CCSB human ORFeome resource version 8.1 (Rual et al. 2005) and were validated by Sanger sequencing. The detail of genes used in this study are shown in **Supplementary Table 1**.

### Construction of destination vectors for functional complementation assays

We constructed Gateway compatible destination vectors for ORF expression in two steps. First, we constructed pDN0520, an expression plasmid with an auxotrophic TRP1 marker by eliminating the DHFR F[1,2] region of pDN0509 (Marchant et al. 2019). This was achieved by inserting a synthesized dsDNA fragment dsOPE001 to the SpeI (New England Biolabs, R3133S) and I-SceI (New England Biolabs, R0694S) digested pDN0509 backbone, by ligation. Ligation reactions were performed using T4 DNA ligase (New England Biolabs, M0202S) following the manufacturer’s instructions. The ligated product was transformed to One Shot™ ccdB Survival™ 2 T1^R^ Competent Cells (Invitrogen, A10460), and selected on LB+Amp plates. The construct was validated using restriction enzyme digestion, followed by Sanger sequencing. We then replaced the backbone containing the marker with CloNAT and Hygromycin resistance to create pDN0603 and pDN0604, respectively. This was achieved by first digesting pDN0520 with I-SceI (New England Biolabs, R0694S), I-CeuI (New England Biolabs, R0699S), and SapI (New England Biolabs, R0569S) to obtain the ADH1pr-Gateway cassette. SapI was used to fragment the backbone, unnecessary for subsequent cloning. The insert region was then ligated to pKB11 and pKB12 (See Construction of destination vectors for DHFR-PCA assays) which were also treated with I-SceI and I-CeuI, giving the backbones with CloNAT and Hygromycin resistance cassettes, respectively. The backbones of pKB11 and pKB12 were treated with calf intestinal alkaline phosphatase (CIP) (New England Biolabs, M0525S) to prevent self ligation. Ligation reactions were performed using T4 DNA ligase (New England Biolabs, M0202S) following the manufacturer’s instructions. These products were transformed to chemically competent *E. coli* strain DB3.1 and selected on LB+Amp plates. The constructs were validated using restriction enzyme digestion, followed by Sanger sequencing.

### Strain preparation for functional complementation assay

We individually subcloned *HRR25* orthologs to pDN0603 (pDEST-ADH1pr Nat^R^) and pDN0604 (pDEST-ADH1pr Hyg^R^) by Gateway LR cloning (Invitrogen, 11791), following the manufacturer’s instructions. Expression vectors in pDN0603 and pDN0604 backbones were used to transform strains GS007 and GS009, respectively. Successful transformants were selected on YPD plates with appropriate antibiotics for selection.

### Functional complementation assay by spot assay

We mated the GS007 and GS009 strains with vehicle and *S. cerevisiae HRR25* plasmids. Diploids were selected on YPD+NAT+HPH plates. We cultured haploid (GS007 or GS009) and diploid (GS007/GS009) strains carrying plasmids in 3mL of YPD medium containing antibiotics for plasmid selection, for approximately 18 hours at 30 °C with agitation. Spots were plated on YPD or YPD+DOX in 5-fold serial dilutions, starting from OD_600nm_ = 0.5.

### Combinatorial functional complementation screening

The 52 haploid strains (26 baits and 26 preys) with plasmids were used to perform the complementation screening on a robotically manipulated pin tool (BM5-SC1, S&P Robotics Inc.). First, the preys (GS009 strains with pDN0604-ortholog constructs) were cherry-picked from 96-well plates to omnitrays in a 384 format on YPD+HPH solid media. Plates were then combined into the 1536 format on YPD+HPH plates. The prey array comprised a border of control strains of two rows and two columns of BY4743, and 26 preys (25 *HRR25* orthologs and vehicle control) each with 9 or more replicates on random assigned positions. The presence of border strain and the random position assignment of preys were performed to minimize the effect of plate position. Second, baits (GS007 strains with pDN0603-ortholog construct) were grown in 10 mL of liquid YPD+NAT, and used to create a ‘lawn’, which we prepared by transferring the overnight culture of bait strains on solid YPD+NAT omnitrays, and removed excess medium after letting the cells settle by waiting for approximately 3 minutes. The plates were dried and incubated at 30 °C for 48 hours. They were then used to perform the mating in 1536 array format. Mating of baits with preys were done on YPD followed by incubation at 30 °C for 48 h. Two diploid selection steps were performed by replicating the plates on YPD+NAT+HPH and incubating them for 48 h at 30 °C. Pictures were taken at the end of the second diploid selection for quality control. All images were acquired with a EOS Rebel T5i camera (Canon). Finally, the diploid cells were replicated to YPD and YPD+DOX media for selection. The plates were incubated for 2 days as first round selection in a spImager custom robotic platform (S&P Robotics Inc.). Cells were replicated for two more rounds on each medium to reduce background for selection and incubated in the same manner. Pictures were taken every 2 hours for the second and third selection for 3 days. The acquired images were processed to obtain colony areas for each time point using pyphe version 0.981 using the pyphe-quantify function (Kamrad et al. 2020). The area under curve (AUC) was computed from the growth curve drawn using colony areas using the trapz function of the numpy library version 1.24. Complementation scores were computed by taking the ratio of AUC values between non-selection (YPD) and selection (YPD+DOX). The AUC values and complementation scores for all constructs are shown in **Supplementary Table 4**.

### Functional complementation assay of chimeric orthologs

The chimeric orthologs were constructed from expression vectors of orthologs in the pDN0603 backbone, namely pDN0603-HRR25 (*S. cerevisiae*), pDN0603-NCAS0A01090 (a) (*N. castellii*), and pDN0603-NCAS0D03820 (B) (*N. castellii*). We PCR amplified the kinase region (corresponding positions 1-290 in *S. cerevisiae HRR25*) of each construct with the backbone region, to produce each construct without the central/tail regions of the ortholog. Amplification was performed using Kapa HiFi Hotstart Polymerase (Roche, 07958897001). The PCR reaction, in a final volume of 25 µL, consisted of 1X reaction buffer, 300 µM of dNTPs, 300 nM of each forward and reverse primer, 0.5 unit of polymerase, and 4 ng of template DNA. Thermal cycling conditions were (i) 95 °C for 5 min; (ii) 30 cycles of 98 °C for 20 s, 60 °C for 15 s, and 72 °C for 1 min; (iii) 72 °C for 1:30 min; and (iv) 12 °C forever. We PCR amplified the central/tail regions of these orthologs with homology to other kinase domains and the backbone. Amplification was also performed using Kapa HiFi Hotstart Polymerase (Roche, 07958897001). The PCR reaction, in a final volume of 25 µL, consisted of 1X reaction buffer, 300 µM of dNTPs, 300 nM of each forward and reverse primer, 0.5 unit of polymerase, and 4 ng of template DNA. Thermal cycling conditions were (i) 95 °C for 5 min; (ii) 30 cycles of 98 °C for 20 s, 60 °C for 15 s, and 72 °C for 10 min; (iii) 72 °C for 15 min; and (iv) 12 °C forever. To construct each of the desired chimeric orthologs, the backbone (with the kinase domain) and the fragment containing central/tail regions were subjected to *in vitro* DNA assembly (Gibson 2009). The resulting products were transformed to chemically competent *E. coli* strain DH5α and selected on LB+Amp plates. Constructs were validated by Sanger sequencing of the inserted region. Primers used to amplify each fragment are listed in **Supplementary Table 3**. The plasmids encoding chimeric orthologs together with controls, were transformed to GS007 using standard lithium acetate transformation, and selected on YPD+NAT. The spot assay was performed as described in *Functional complementation assay by spot assay*.

### Molecular visualization

The protein crystal structure of Hrr25p (PDB: 5CYZ) (Ye et al. 2016) was visualized using ChimeraX (version 1.5) (Pettersen et al. 2021).

### Plasmid construction for localization assay

We constructed Gateway compatible destination vectors with C-terminal GFP fusion. This plasmid, pDN0617 (pDEST-Gateway-GFP Nat^R^), was constructed by cloning the mEGFP to pDN0603 (pDEST-Gateway Nat^R^).

We first constructed pUG27-mEGFP-ENO1t-SpHIS5 by inserting mEGFP and ENO1 terminator fragments to pUG27 (Gueldener et al. 2002) using *in vitro* DNA assembly (Gibson et al. 2009). All three fragments were PCR amplified using the Kapa HiFi Hotstart Polymerase (Roche, 07958897001), and treated with 1 µL (20 units) of DpnI (New England Biolabs, R0176L) for 1 h at 37°C before being purified on magnetic beads. The backbone fragment was amplified from pUG27 using primers CLOP161-F8 and CLOP161-F9, with the thermal cycling conditions as (i) 95 °C for 5 min; (ii) 35 cycles of 98 °C for 20 s, 68°C for 15 s, and 72 °C for 2 min; (iii) 72 °C for 3 min; and (iv) 12 °C forever. The mEGFP fragment was amplified from the plasmid F65V-34a-mEGFP using primers CLOP161-F10 and CLOP161-F12. The ENO1 terminator fragment was amplified from pYTK051 (Addgene Kit #1000000061 position E3) (Lee et al. 2015) using primers CLOP161-G1 and CLOP161-G2. Both the mEGFP and ENO1 terminator fragments were amplified using the following thermal cycling condition (i) 95 °C for 5 min; (ii) 5 cycles of 98 °C for 20 s, 60 °C for 15 s, and 72 °C for 30 s; (iii) 30 cycles of 98 °C for 20 s, 72 °C for 45 s; (iv) 72 °C for 3 min; and (v) 12 °C forever. All PCR reactions, in a final volume of 25 µL, consisted of 1X reaction buffer, 300 µM of dNTPs, 300 nM of each forward and reverse primer, 0.3 unit of polymerase, and 10 ng of template DNA. Five microliters of the *in vitro* DNA assembly reaction were transformed into MC1061, and plated on LB+Amp. Correct construction of the plasmid was confirmed by Sanger sequencing.

We then constructed pDN0617 by cloning the mEGFP fragment of pUG27-mEGFP-ENO1t-SpHIS5 to pDN0603. The mEGFP fragment with homology to the pDN0603 backbone was amplified from pUG27-mEGFP-ENO1t-SpHIS5 using primers CLOP286-A11 and CLOP286-B11. Amplification was performed using Kapa HiFi Hotstart Polymerase (Roche, 07958897001). The PCR reaction, in a final volume of 25 µL, consisted of 1X reaction buffer, 300 µM of dNTPs, 300 nM of each forward and reverse primer, 0.5 unit of polymerase, and 0.4 ng of template DNA. Thermal cycling conditions were (i) 95 °C for 5 min; (ii) 30 cycles of 98 °C for 20 s, 60 °C for 15 s, and 72°C for 1 min; (iii) 72 °C for 1:30 min; and (iv) 12 °C forever. The mEGFP insert was column purified after DpnI (New England Biolabs, R0176S) treatment, and subjected to *in vitro* DNA assembly (Gibson et al. 2009) with SpeI-HF (New England Biolabs, R3133S) digested pDN0603 as backbone. This reaction was transformed to chemically competent *E. coli* strain DB3.1 and selected on LB+Amp plates. The construct was validated using restriction enzyme digestion, followed by Sanger sequencing.

### Strain construction for Western blotting and localization assay

We individually subcloned *HRR25* orthologs to pDN0617 (pDEST-Gateway-GFP Nat^R^) by Gateway LR cloning (Invitrogen, 11791), following the manufacturer’s instructions. Expression vectors in pDN0617 backbones were used to transform BY4741. Successful transformants were selected on YPD+NAT plates.

### Western blotting

First, BY4741 strains carrying plasmid with each ortholog tagged with GFP were grown overnight in SC complete medium+NAT. The next morning, yeast cells were diluted to 0.15 OD_600nm_ and grown for 6 hours to reach exponential phase (1 OD_600nm_). From this culture, 12.5 OD_600nm_ units of cells were collected and freezed at −80°C. Each pellet was resuspended in 250 µL lysis buffer (Complete Mini Protease inhibitor cocktail, Millipore-Sigma 11836153001) and glass beads were added to the suspension. The mixture was vortexed for 5 min on a Turbomix. After vortexing, 25 µL of 10% SDS was added to each tube and then boiled for 10 min. Tubes were then centrifuged at 16,000×g for 5 min to clear the supernatant. Samples for SDS-PAGE gel were prepared by mixing 17.5 µL cell extract, 2.5 µL DTT 1M and 5 µL LB 5X (250mM Tris-Cl pH 6.8, 10% SDS, 0.5% Bromophenol Blue, 50% glycerol). Samples were run on 8% SDS-PAGE gel for 1 hour at 175V (Tetra cell, BioRad). Following migration, proteins were transferred to a nitrocellulose membrane for 75 min at 0.8 mA/cm^2^. Once on the nitrocellulose membrane, proteins were stained with Ponceau Red to check proper and uniform loading/transfer (Romero-Calvo et al. 2010). Following the imaging of the Ponceau Red staining, the membranes were blocked overnight in the blocking solution (Intercept^®^ (PBS) Blocking Buffer, 927-70003, Mandel Scientific) at room temperature with agitation. The next morning, membranes were soaked in blocking solution +0.02% Tween 20 with 0.15 ug/ml GFP antibody (Millipore-Sigma 11814460001) for 30 min. After the primary antibody incubation, the membranes were washed 3 times each for 5 min with PBS+T (10 mM sodium phosphate dibasic, 135 mM NaCl, 2 mM KCl, 1.5 mM monopotassium phosphate, 0.1% Tween 20). For the secondary antibody, membranes were again soaked in blocking solution +0.02% Tween 20 with 0.075 µg/ml Anti-Mouse 800 (LIC-926-32210, Mandel Scientific) for 30 min. Membranes were washed another 3 times each for 5 min and then imaged with an Odyssey Fc instrument (Licor, Mandel Scientific) in 700 and 800 channels.

### Construction of destination vectors for DHFR-PCA assays

We constructed gateway compatible DHFR-PCA plasmids pKB11 (pDEST-DHFR F[1,2] C-term Nat^R^), pKB12 (pDEST-DHFR F[3] C-term Hyg^R^), and pDN0605 (pDEST-DHFR F[1,2] N-term Nat^R^) containing Nat^R^ marker, tagging the genes of interest in the C and N-terminus, respectively. We first constructed pKB11 by replacing the TRP1 auxotrophic cassette of pDEST-DHFR F[1,2]-C (TRP1) (Addgene #177795) (Marchant et al. 2019) to the Nat^R^ marker. We amplified the Nat^R^ cassette using primers CLOP148-G1 and CLOP148-G2 from pAG25 (Euroscarf, P30104). The backbone region without the auxotrophic marker was prepared by PCR amplification of pDEST-DHFR F[1,2]-C (TRP1), using primers CLOP157-H1 and CLOP157-H2. These insert and backbone fragments were subjected to *in vitro* DNA assembly to create pKB11. Correct construction of pKB11 was confirmed by Sanger sequencing. In a similar way, pKB12 was constructed by replacing the LEU2 auxotrophic cassette of pDEST-DHFR F[3]-C (LEU2) (Addgene #177796) (Marchant et al. 2019) to the Hyg^R^ marker, using pAG32 (Euroscarf, P30106). The N-terminus fusion plasmid pDN0605 was constructed using pDEST-DHFR F[1,2]-N (TRP1) (Addgene #177797) (Evans-Yamamoto et al. 2022), and replacing the backbone region containing the selection marker with that of pKB11. This was achieved by restriction enzyme digestion followed by ligation. Both pDN0605 and pKB11 were digested with I-SceI (New England Biolabs, R0694S) and I-CeuI (New England Biolabs, R0699S) homing endonucleases to obtain the ADH1pr-DHFR F[1,2]-Gateway cassette and backbone, respectively. The digested product of pKB11 was treated with CIP (New England Biolabs, M0525S) to prevent self ligation, before ligation.

### DHFR-PCA assay

We individually subcloned *HRR25* orthologs to pKB11 (pDEST-DHFR F[1,2] C-term Nat^R^) and pDN0605 (pDEST-DHFR F[1,2] N-term Nat^R^) by LR cloning (Invitrogen, 11791), following the manufacturer’s instructions. The plasmids were purified and used to transform BY4741 strains using standard lithium acetate transformation protocol. The pKB11 and pDN0605 constructs were transformed into BY4741, and selected on YPD+NAT plates for successful transformants. We used these strains to test interactions against the DHFR F[3] miniarray, in the BY472 background. The DHFR F[3] miniarray is a collection of strains which genomically fused 1,172 genes to the DHFR F[3] reporter in *S. cerevisiae*, which have been reported to have at least one PPI using DHFR-PCA (Tarassov et al. 2008; Rochette et al. 2015; Diss et al. 2017). The full list of proteins in the DHFR F[3] miniarray used in this assay is shown in **Supplementary Table 5**. The DHFR-PCA screening was performed on a robotically manipulated pin tool (BM5-SC1, S&P Robotics Inc.) as previously described (Rochette et al. 2015).

First, the preys (DHFR F[3] miniarray collection) were grown on omnitrays in a 384 format on YPD+HPH solid media. Plates were then combined into the 1536 format on YPD+HPH plates. The prey array comprised a border of control strains of two rows and two columns corresponding to the PPI between LSM8-DHFR F[1,2] and CDC39-DHFR F[3]. The presence of border strain and the random position assignment of preys were performed to minimize the effect of plate position. Second, baits (strains each carrying one of the *HRR25* orthologs) were grown in 10 mL of liquid YPD+NAT, and used to create a ‘lawn’, which we prepared by transferring the overnight culture of bait strains on solid YPD+NAT omnitrays, and removed excess medium after letting the cells settle by waiting for approximately 3 minutes. The plates were dried and incubated at 30 °C for 48 hours. They were then used to perform the mating in 1536 array format. Mating of DHFR F[1,2] baits with DHFR F[3] preys was done in two independent replicates on YPD followed by incubation at 30 °C for 48 h. Two diploid selection steps were performed by replicating the plates on YPD+NAT+HPH and incubating them for 48 h at 30 °C. Pictures were taken at the end of the second diploid selection for quality control. All images were acquired with a EOS Rebel T5i camera (Canon). Finally, the diploid cells were replicated on omnitrays containing solid PCA selection media and incubated for 4 days as a first selection step in a spImager custom robotic platform (S&P Robotics Inc.). Cells were replicated for a second PCA selection step and incubated in the same manner. Pictures were taken every 2 hours for the second PCA selection for 4 days. The acquired images were processed to obtain colony areas for each time point using pyphe version 0.981 using the pyphe-quantify function (Kamrad et al. 2020). The area under curve (AUC) was computed from the growth curve drawn using colony areas using the trapz function of the numpy library version 1.24. Colonies were quality filtered based on the colony area of the diploid selection plate, where colonies with Log_2_ transformed area below 10 were excluded for PPI evaluation. We set a threshold to call positive interactions at AUC = 1.0 × 10^4^, based on the distribution of AUCs among tested pairs. We note that, among the two tagging orientations (N and C-terminal tagging) tested for *HRR25* orthologs, only the C-terminal tagged constructs showed interactions. Subsequent analyses were performed on the data generated from the C-terminal tagged constructs only. The PPI score data generated from the C-terminal tagged constructs are shown in **Supplementary Table 6**.

### GO enrichment analysis of PPI partners

PPI partners for each ortholog were classified according to their gene ontology (GO) term as implemented in the enrichGO function of the R package clusterProfiler (Yu et al. 2012), using gene sets of cellular components in the *S. cerevisiae* database. P-values were adjusted using the Benjamini and Hochberg method. GO terms with the same description were combined and represented with the lowest adjusted p-value for visualization. The data regarding the GO enrichment analysis are shown in Supplementary Table 7.

### SH3 domain motif analysis

To evaluate the SH3 binding motifs, we used position weight matrices (PWMs) for each SH3-domain constructed from quantitative motif binding assays (Tonikian et al. 2009; Jain and Bader 2016). We evaluated the occurrences of PWMs and its similarity within each ortholog, using matrix similarity score (MSS). We first created a null distribution of MSS by creating a randomized protein set with the same length as the *HRR25* orthologs with amino acid frequencies of the *S. cerevisiae* proteome. For each ortholog-PWM combination, the highest value was adopted to represent the motif score, unless multiple values were observed above the 95^th^ percentile of the random distribution (MSS = 0.64). The full list of MMS values for each ortholog-PWM combinations are shown in **Supplementary Table 8**.

### Microscopy experiments

Cells expressing the mEGFP fusion orthologs were grown until early log phase, by first culturing overnight in 5 mL of SC complete medium+NAT, and diluted 1:50 in fresh medium 3 hours prior to imaging. The cells were then seeded on 0.05 mg/mL concanavalin A (Millipore Sigma) coated 96 well glass bottom plate (Cellvis, P96-1.5H-N). Image acquisition was performed using a Nikon Eclipse TE2000-U inverted microscope equipped with a Perkin Elmer UltraVIEW confocal spinning disk unit, a Plan Apochromat DIC H 100×/1.4 oil objective (Nikon), and a Hamamatsu Orca Flash 4.0 LT + camera. Imaging was done at 30 °C in an environmental chamber. The software NIS-Elements (Nikon) was used for image capture. Cells were excited with a 488 nm laser set at 60% intensity and emission was filtered with a 530/630 nm filter to acquire specifically GFP fluorescence. For each field, one brightfield and 5 Z-stacked (2 µm apart) fluorescence images were acquired.

### Gene expression analysis

Gene expression datasets for *S. cerevisiae*, *K. lactis*, and *N. castellii* were downloaded from the Gene Expression Omnibus (GEO) with the accession IDs GSE36599 (Xue-Franzén et al. 2013), GSE22198 (Tsankov et al. 2010), and GSE90041 (Szachnowski et al. 2019). In order to compare ortholog gene expression levels among different datasets, 1:1:1 orthologs for the three species were identified based on the YGOB database annotation (Byrne and Wolfe 2005). Gene expression levels were Log_2_ transformed, and normalized by subtracting the mean of Log_2_ transformed expression level of 1:1:1 orthologs. The *N. castellii* gene pairs which were maintained from WGD were evaluated for expression imbalance among ohnologs. Gene pair information was extracted based on the YGOB database annotation (Byrne and Wolfe 2005), and raw read counts were used for gene expression levels. We note that the position information does not reflect the loci which the ohnologs are encoded. Expression imbalance was computed by taking the ratio between the raw read count of ohnolog pairs.

### Essentiality assay in *N. castellii*

Gene deletion in *N. catselli* was performed as described in (Karademir Andersson et al. 2016). First, an antibiotic marker (HPHNT1 or NATNT2) was fused by PCR with 500 bp homology with the promoter and terminator of each ohnolog (NCAS0A01090 and NCAS0D03820). This was achieved by fusion PCR of three PCR fragments, each containing the antibiotic marker (amplified from plasmids pFA6a-natNT2 and pFA6a-hphNT1 (Janke et al. 2004) with homology to 5′ and 3′ regions of the target gene), the 500 bp 5′ region of the target gene, and the 500 bp 3′ region of the target gene. Primers used to amplify each fragment are listed in **Supplementary Table 3**.

Genomic integration of the PCR product was performed in the diploid strain obtained by crossing *N. castellii* strains YMC7 and YMC8 (Karademir Andersson et al. 2016). The cells were grown overnight in YPD at room temperature to reach OD_600nm_ around 0.8. The cells were washed once with sterile water and a second time with transformation buffer I (0.1M lithium acetate, 10mM Tris–HCI, 1 mM EDTA, pH 7.5). Following the second wash, the pellet was resuspended in 250 µL transformation buffer I. The transformation reaction was set up using 8 µL of the PCR fusion, 2 µL DNA carrier 10mg/ml with 50 µL competent cells and 300 µL transformation buffer II (40% PEG 3350, 1× transformation buffer I). The reaction was incubated 30 min at room temperature followed by a 30 min heat shock at 30°C. After the heat shock, the cells were washed twice with 1 ml sterile water before being resuspended in 100 µL of sterile water for plating on YPD with the appropriate antibiotics (HPH 200 µg/mL and NAT 50µg/mL). After 3 days at room temperature, isolated colonies were tested by colony PCR (Lõoke et al. 2011).

Once deletion was confirmed by PCR, the cells were transferred to sporulation media (Potassium acetate 0.3%, Yeast Extract 0.1%, Glucose 0.05%, Amino-acid supplement powder (2g histidine, 10g leucine, 2g lysine, 2g uracil) 0.01%, Agar 2%) for tetrad dissection. After 5 days on sporulation media, cells were resuspended in zymolyase 20T solution (200 µg/mL) for 20 min at room temperature. Following the zymolyase treatment, cells were spun down for 15 sec at 16,000 × g and resuspended in 200 µL sorbitol 1M. Treated cells were then placed on a level YPD plate to perform tetrad dissection with the SporePlay (Singer Instrument). The dissection plates were incubated at room temperature for 5 days to make sure we didn’t underestimate growth. Once the spores were grown enough, each spore (by keeping tetrad association) were resuspended in 200 µL sterile water and 5 µL was spotted on YPD, YPD+NAT (50 µg/mL), YPD+HPH (200 µg/mL) and YPD+G418 (75 µg/mL) for viability evaluation.

## Supplementary Material

**Supplementary Table 1.** List of genes in this study.

**Supplementary Table 2.** Biological resources used in this study.

**Supplementary Table 3.** List of oligonucleotides used in this study.

**Supplementary Table 4.** Data from the functional complementation assay.

**Supplementary Table 5.** List of strains contained in the DHFR F[3] miniarray.

**Supplementary Table 6.** Data from the PPI screening.

**Supplementary Table 7.** GO enrichment values from detected PPIs.

**Supplementary Table 8.** Motif match score (MMS) for each ortholog.

## Data Availability Statement

The data underlying this article are available in the article and in its online supplementary material. Biological resources will be available though Addgene or by request to the corresponding author. The computer codes and control files, defining the settings for the codeml program execution, used in this study are available at GitHub (https://github.com/Landrylab/Evans-Yamamoto_et_al_2023, last accessed September 26^th^, 2023).

## Author contributions

DEY and CRL designed the study. CRL supervised the work. DEY, GS, AKD, and IGA prepared the material for assays using *S. cerevisiae*. DEY, GS, and AKD performed the complementation and PPI assays. AKD operated the S&P robotic platform for complementation and PPI assays. SP performed the imaging experiments. AKD performed the Western blot analysis and experiments using *N. castellii*. DEY performed all analysis with support from DB and input from CRL. DEY and CRL wrote the initial draft of the manuscript. DEY, AKD, GS, SP, DB, and CRL contributed to revising, reading and approving the final version of the manuscript.

## Acknowledgments

The *N. castellii* strain CBS 4309 was a kind gift from Dr. Kenneth H Wolfe. The *N. castellii* strain YMC7 and YMC8 were a kind gift from Dr. Marita Cohn. The F65V-34a-mEGFP plasmid was a kind gift from Dr. Michael Springer. Molecular graphics was visualized with UCSF ChimeraX, developed by the Resource for Biocomputing, Visualization, and Informatics at the University of California, San Francisco, with support from National Institutes of Health R01-GM129325 and the Office of Cyber Infrastructure and Computational Biology, National Institute of Allergy and Infectious Diseases. This project was partially funded by a Canadian Institutes of Health Research Foundation grant number 387697 and a NSERC discovery grant to CRL. This work was also supported by the JSPS DC1 fellowship (to DEY), TTCK fellowship (to DEY), Watanabe Foundation International Scholarship (to DEY), and MITACS fellowship (to GS). CRL holds the Canada Research Chair in Cellular Systems and Synthetic Biology.

---

**Supplementary Figure 1.**
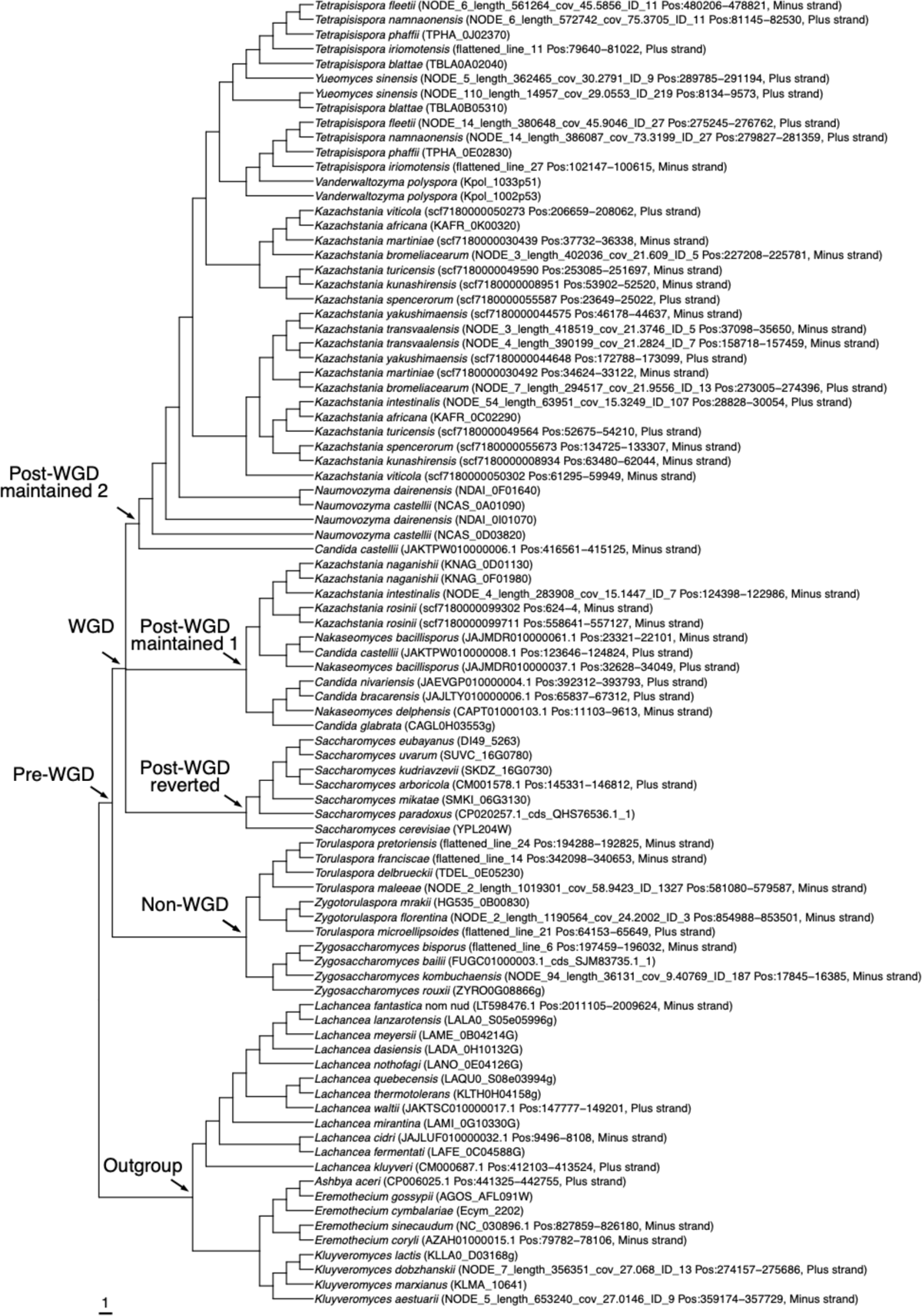
Reconciliated gene tree of *HRR25* orthologs.

**Supplementary Figure 2.**
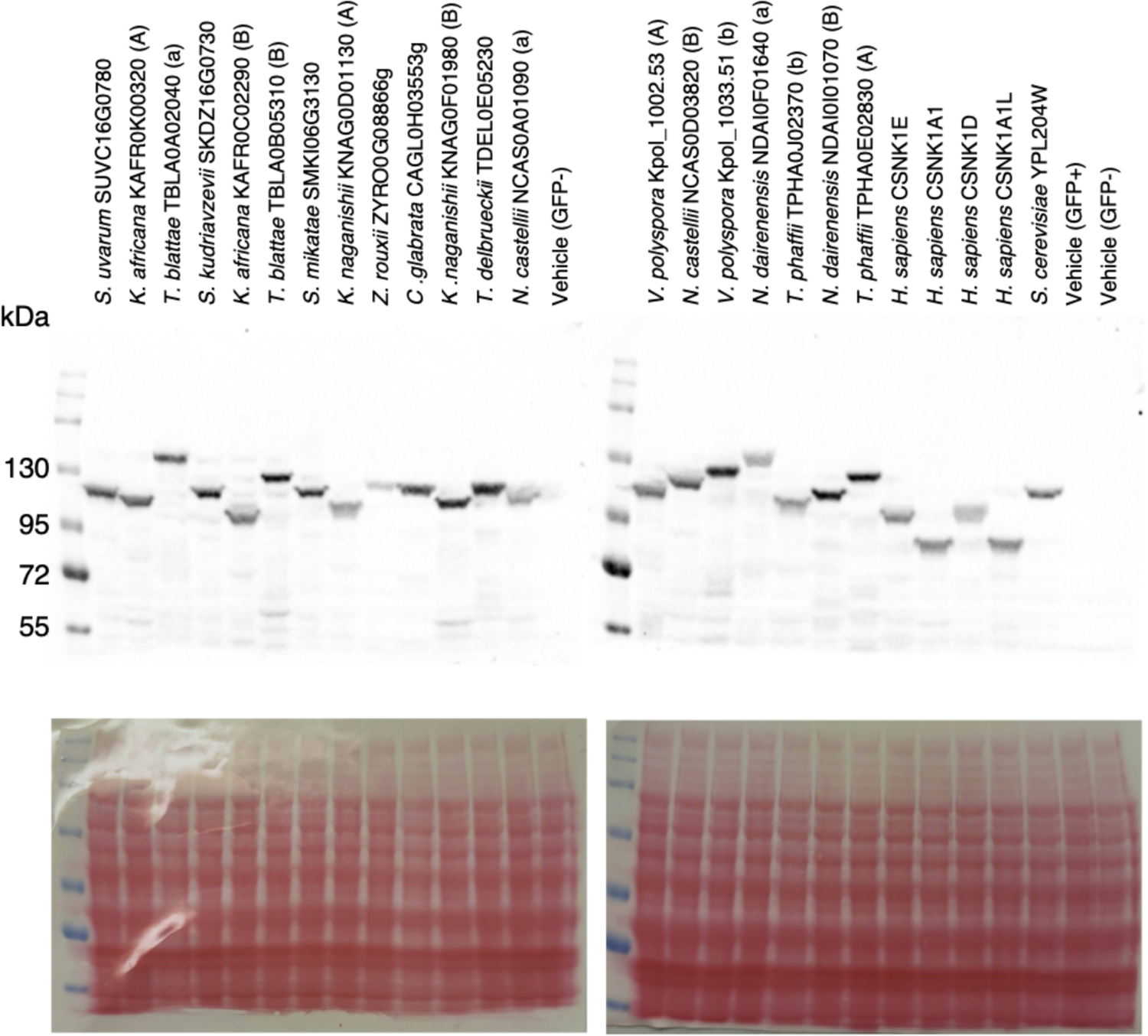
*HRR25* orthologs are expressed in *S. cerevisiae*. (**Top**) Protein level expression of each ortholog, expressed with C-terminal mEGFP fusion. Western blot using an anti-GFP antibody (Millipore-Sigma 11814460001). Variation in migration is expected given that the length of the PQ-rich tail region is variable among orthologs. (**Bottom**) Images for Ponceau staining of membrane prior to antibody incubation, as loading control (Romero-Calvo et al. 2010). Vehicle (GFP+): Pre-LR vehicle control encoding mEGFP (pDN0617) with no GFP expression due to absence of promoter region in proximity. Vehicle (GFP−): Pre-LR vehicle control, which does not encode mEGFP (pDN0603).

**Supplementary Figure 3.**
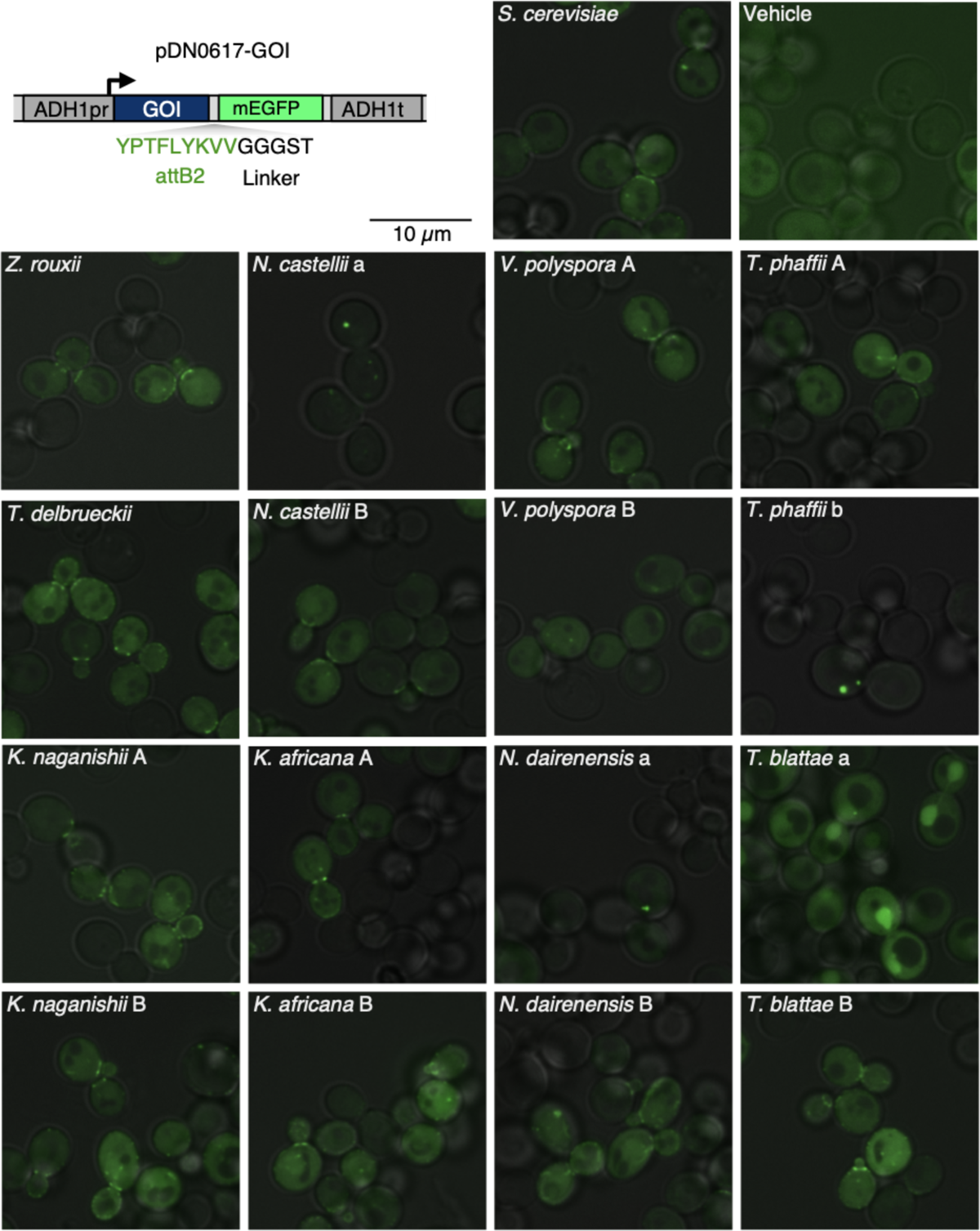
Localization of mEGFP-fused *HRR25* orthologs in *S.cerevisiae*.

